# Neuroinflammation and metabolic dysfunction in POLG-related mitochondrial epilepsy

**DOI:** 10.64898/2026.08.12.744403

**Authors:** Laura A. Smith, Megan Wilson, Elsaid Mohamed Elsaid, Pawel Palmowski, Zirong Jiang, Laura Aryeetey, Conor Holly, Jenna Dickin, Maxwell Abbey, Anna L. Smith, Robert W. Taylor, Omar Hikmat, Charalampos Tzoulis, Gavin Hudson, Daniel Erskine, Robert McFarland

## Abstract

Super-refractory status epilepticus is a common neurological manifestation of mitochondrial disease caused by bi-allelic pathogenic variants in *POLG*. Epilepsy in POLG-related disease typically presents with an explosive onset of status epilepticus, often from an occipital focus, and is associated with extensive neurodegeneration. The neuropathological mechanisms underlying POLG-related mitochondrial epilepsy remain poorly understood, however, neuroinflammation and glial dysfunction are hypothesised to play a significant role. In this study, we performed a neuropathological and proteomic investigation of post-mortem brain tissues from 12 patients with POLG-related mitochondrial epilepsy (age range: 3 – 28 years) and matched control cases. Given that the primary visual cortex is prominently involved in this epileptic disorder, occipital cortical tissues (Brodmann area 17) were compared to frontal cortical tissues (Brodmann area 9). Liquid chromatography-mass spectrometry (LC-MS/MS) analysis identified a distinct immunometabolic signature in the occipital cortex, and to a lesser extent in the frontal cortex, in POLG-related epilepsy. This was characterised by decreased abundance of mitochondrial proteins coupled to an increased expression of innate immune and inflammatory proteins, consistent with neuroinflammation. To validate these observations, we confirmed an increased density of cells immunoreactive for acute phase proteins (C-reactive protein, osteopontin and serpin A3), immune co-receptors (CD14 and HLA-DR), the inflammatory glycoprotein YKL40, the cytokine TNF-alpha, and mitochondrial translocator protein (TSPO). We also demonstrate a decreased expression of mitochondrial oxidative phosphorylation (OXPHOS) subunits within POLG patient microglia, indicative of mitochondrial dysfunction. Finally, we show enrichment of mitochondrial OXPHOS and interneuron proteins in the control primary visual cortex compared with the frontal cortex, which may underlie the selective regional vulnerability observed in POLG-related mitochondrial disease. Overall, these findings provide strong neuropathological evidence implicating neuroinflammation and glial dysfunction in POLG-related epilepsy.

## Introduction

Bi-allelic pathogenic variants in the mitochondrial DNA polymerase gamma gene (*POLG*) are a leading cause of epilepsy in patients with primary mitochondrial disease [41, 65]. POLG-related epilepsy typically presents with an explosive onset of status epilepticus, is super-refractory to anti-seizure medications, and shows a predilection for the primary visual cortex [3, 9]. Whilst this epilepsy can present at any age, it is particularly relentless in early-and juvenile-onset POLG-related disease, including young adults [17], and likely underpins the accelerated loss of neurological function in patients.

The precise pathophysiological mechanisms leading to the development of POLG-related epilepsy remain unknown. However, bi-allelic pathogenic *POLG* variants are known to impair mitochondrial DNA (mtDNA) replication, leading to a depletion of mtDNA and subsequent disruption of the oxidative phosphorylation (OXPHOS) system [33]. Since mitochondria have a role in regulating innate immune responses [64], and are important sources of damaged-associated molecular patterns (DAMPs) including reactive oxygen species (ROS) and mtDNA [63], mitochondrial dysfunction due to pathogenic *POLG* variants is hypothesised to disrupt innate immune signalling events.

There is mounting clinical and experimental evidence supporting the dysregulation of the innate immune system in POLG-related disease. This includes reports that the onset of status epilepticus follows an infectious period in nearly one third of patients with POLG-related disease [16]. Furthermore, elevated levels of interferon-related genes including *IFI27* have been observed in blood samples from paediatric patients with mitochondrial disease, including patients with mtDNA maintenance defects [23], in addition to increased cytokines in cerebral spinal fluid samples from an infant with Alpers’ syndrome [14]. Moreover, the pathogenic *POLG* variant p.Trp748Ser (NM_002693.3: c.2243G>C) has been linked with immunodeficiencies and an aberrant anti-viral response [20], and a mouse model harbouring a pathogenic *Polg* variant (*Polg^D257A^*) demonstrates a hyperinflammatory phenotype characterised by an elevated type I interferon response [31, 59].

Although there is evidence providing a direct link between defective POLG and dysfunction and/or heightened activation of the innate immune system in POLG-related disease [20, 59], the neuroinflammatory mechanisms involved in POLG-related epilepsy remain poorly characterised. However, activated microglia (the resident immune cells of the brain), in addition to reactive astrocytes, are known to promote neuroinflammation and neuronal hyperexcitability through the release of pro-inflammatory cytokines [42]. Therefore, mitochondrial dysfunction may act as a direct stimulus of innate immune activation [64], and play a causal role in the development of POLG-related epilepsy. In support of this, previous neuropathological studies examining *post-mortem* brain tissues from patients with POLG-related disease revealed an accumulation of abnormal reactive astrocytes and diffuse microglial activation in occipital cortical tissues [6, 48, 58].

To elucidate the neuroimmune and neurodegenerative mechanisms underpinning epilepsy in patients with POLG-related disease, we performed a discovery proteomics and neuropathological study using *post-mortem* cortical tissues from patients with early-onset and juvenile to adult-onset POLG-related disease. Here, we provide evidence of an immunometabolic proteome characterised by a decreased abundance of mitochondrial proteins coupled to an increased abundance of innate immune and inflammatory proteins. We also present data highlighting an enriched metabolic and inhibitory interneuron proteome in control primary visual cortex tissues which could partly explain the vulnerability of this brain region to mitochondrial dysfunction in POLG-related disease.

## Materials and methods

### Patient and control post-mortem tissue cohort

Human post-mortem cortical tissues were obtained from brain banks located in the United Kingdom (Newcastle Brain Tissue Resource, The Oxford Brain Bank, Edinburgh Brain Tissue Resource and University Hospital Southampton), Norway (Neuro-SysMed Center Brain Bank, Haukeland University Hospital, Bergen) and the United States (NIH NeuroBioBank, University of Maryland). Tissues were obtained from 12 patients with mitochondrial epilepsy due to bi-allelic pathogenic *POLG* variants (age range: 3 years – 28 years; **Table 1** and **Supplementary Table 1**) and were compared to control cases who died from non-neurological causes and had no significant neuropathological abnormalities identified at post-mortem examination (**Supplementary Table 2 – 3**). Ethical approval was obtained from the individual brain banks.

**Table 1.** POLG-related epilepsy patient cohort.

| Case ID | Age of onset | Age at death | Sex | Bi-allelic POLG variants | Epilepsy | Stroke-like episode | Cortical visual symptoms | Developmental delay | Liver dysfunction / failure | EEG recordings | Neuroimaging lesions | References |
| --- | --- | --- | --- | --- | --- | --- | --- | --- | --- | --- | --- | --- |
| Pt.01 | 2 y | 3 y | M | p.[Cys418Arg]/p.[Ala467Thr] | + | + | x | + | x | Epileptic activity | Occipital, frontal, parietal, thalamus | [50] |
| Pt.02 | 3 y | 3 y | F | p.[Ala467Thr]/p.[Arg852Cys] | + | x | x | + | x | Not available | Occipital, thalamus | Unpublished |
| Pt.03 | 2 y | 11 y | M | p.[Ala467Thr]/p.[Gly848Ser] | + | x | + | + | + | Slow posterior background rhythm over both occipital lobes; frequent generalised polymorphic delta activity | Not available | [48-50] |
| Pt.04 | 1 y | 13 y | M | p.[Leu424fs]/p.[Ala467Thr] | + | x | x | + | x | Abnormal background rhythm; multi-focal epileptiform activity; clinical myoclonus without EEG correlate | Not available | Unpublished |
| Pt.05 | 2 y | 13 y | M | p.[Trp748Ser]/p.[Trp748Ser] | + | + | + | + | + | Multi-focal epileptiform activity; right and left side parietal occipital activity | Occipital, thalamus, cerebellum | [58] |
| Pt.06 | 18 y | 23 y | F | p.[Ala467Thr]/p.[Ala467Thr] | + | + | + | x | + | Continuous posterior spike-and-wave activity | Occipital, parietal | [10, 34, 48-50] |
| Pt.07 | 20 y | 24 y | F | p.[Ala467Thr]/p.[Trp748Ser] | + | + | + | x | x | Encephalopathic posterior quadrant spike waves | Occipital, thalamus, parietal, cerebellum | [6, 27-30, 34, 48-50] |
| Pt.08 | 8 y | 24 y | F | p.[Ala467Thr]/p.[Arg597Trp] | + | + | + | x | x | Occipital seizures evident | Occipital | Unpublished |
| Pt.09 | 15 y | 24y | F | p.[Trp748Ser]/p.[Trp748Ser] | + | + | + | – | + | Slow posterior background rhythm over both occipital lobes; epileptiform activities in right occipital lobe | Occipital, thalamus | [58] |
| Pt.10 | 16 y | 24 y | F | p.[Trp748Ser]/p.[Trp748Ser] | + | + | + | – | + | Slow posterior background rhythm over right occipital lobe; epileptiform activities in right temporal/occipital | Occipital, thalamus, pons, cerebellum | [57, 58] |
| Pt.11 | 12 y | 28 y | M | p.[Trp748Ser]/p.[Trp748Ser] | + | + | – | – | + | Slow posterior background rhythm over both occipital/parietal lobes; epileptiform activities in right occipital | Occipital, thalamus | [57, 58] |
| Pt.12 | 16 y | 28 y | F | p.[Ala467Thr]/p.[Trp748Ser] | + | + | + | x | x | Not available | Occipital, frontal, cerebellum | [34, 48-50] |

Abbreviations: y years; m months; M male; F female; FFPE Formalin-fixed paraffin-embedded. POLG Ref\_Seq NM\_002693.3.
Clinical symptoms: + (reported symptom); – (reports that symptom was not present); x (symptom not recorded in clinical-neuropathological reports).
+ FFPE tissues were included in the neuropathological study; + Frozen tissues were included in the proteomics study.

To identify region-specific changes to the primary visual cortex, the predominant site of epileptogenesis in POLG-related epilepsy, occipital cortex tissues (Brodmann area 17/BA17) were compared to frontal cortex tissues (Brodmann Area 9/BA9), which is typically less severely involved in this epileptic disorder. Frozen cortical tissues from 10 patients and 11 control cases were included in a discovery mass spectrometry proteomics study and were matched for age at death (Mann-Whitney test, *P* > 0.05), post-mortem interval (PMI; Mann-Whitney test, *P* > 0.05) and sex (Fisher’s exact test, *P* >0.999; **Supplementary Table 2** and **Supplementary Table 4**). FFPE tissues from 9 patients and 8 control cases were included in a neuropathology study, matched for age at death (Mann-Whitney test, *P* > 0.05) and sex (Fisher’s exact test, *P* >0.999; **Supplementary Table 3** and **Supplementary Table 4**). However, it was not possible to match the PMI between patients and controls for the FFPE tissue cohort (unpaired t test, *P* = 0.0202).

### Sample preparation for untargeted proteomics

Cortical tissue lysates were carefully dissected to remove meninges and underlying white matter, then homogenised in 0.2M triethylammonium bicarbonate (TEAB) buffer with protease inhibitors. The protein concentration of each sample was assessed using a Micro BCA^TM^ Protein Assay Kit (Pierce^TM^). 30µg of total protein was used for processing and trypsin digestion. The total volume of each sample was adjusted to 20µl with S-Trap^TM^ lysis buffer (pH 8.5) composed of 5% sodium dodecyl sulphate (SDS) and 50 mM TEAB. Samples were then reduced in 50mM dithiothreitol (DTT) for 30 minutes at 65°C, alkylated in 100mM iodoacetamide for 30 minutes at room temperature in the dark, and acidified with phosphoric acid to a final concentration of 2.5%. Following this, samples were loaded onto S-Trap^TM^ micro spin columns in 6x volume of S-Trap binding buffer (100mM TEAB in 90% methanol) and centrifuged at 4000 x g for 30 seconds, then washed three times with binding buffer and digested for 8 hours at 37°C with trypsin (Worthington, Lakewood, NJ, US) in 50mM TEAB, at a ratio of 10:1 protein to trypsin. Peptides were eluted in three steps: 50 µL of 50mM TEAB, 50 µL of 0.1% formic acid, and 50 µL of 50% of acetonitrile with 0.1% formic acid. The eluate was dried and reconstituted in 20 µL of 2% acetonitrile with 0.1%.

### Mass spectrometry proteomics

Liquid chromatography-mass spectrometry (LC-MS/MS) was performed on an Ultimate 3000 Rapid Separation LC (RSLC) nano LC system (Thermo Corporation) coupled with the Thermo Scientific™ Orbitrap Fusion Lumos mass spectrometer. 1µg of each sample was loaded onto an AcclaimTM PepMapTM 100 C18 LC Column (5 mm x 0.3 mm, 5 µm particles, 10 nm pores, Thermo Fisher Scientific) at a flow rate of 10 μL⋅min^−1^, at 45 °C. The peptides were separated on an EasySpray C18 column (75 µM x 75 cm, 2 µm particles, Thermo Fisher Scientific) at 45°C, using a 60 minute gradient from 92.5 % A (0.1% formic acid in 3% DMSO) and 8.5% B (0.1% formic acid in 80% acetonitrile 3% DMSO), to 35 % B, at a flow rate of 150 nL min^−1^. Following the separation, peptides were directed to the mass spectrometer through an EasySpray source at the Ion Transfer Tube temperature of 275°C, spray voltage 1600V and analysed using data independent (DIA) acquisition. The mass spectrometry resolution was set to 120,000, with a normalized automatic gain control (AGC) target of 1250%, maximum injection time of 100ms and scan range of 375−1250 mass-to-charge ratio (m/z). Data-independent acquisition (DIA) MS/MS were acquired using 15 windows covering 350-1275 m/z range, at 60000 resolution, AGC target set to 10000%, maximum injection time of 100ms and normalized collision energy (NCE) level of 30%.

The acquired data was analysed in DIA-NN 2.1.0 against the human proteome database (Uniprot UP000005640, accessed January 2024) combined with the common Repository of Adventitious Proteins (cRAP). Search parameters were as follows: fragment m/z 200-1800; enzyme, trypsin with one allowed missed-cleavage; peptide length 7-30 amino acids; precursor m/z 300-1800; precursor charge 2-4; fixed modifications, carbamidomethylation (C); variable modifications, oxidation (M) and acetylation (N-term).

### Proteomic analysis

Proteomics data were analysed using Perseus (1.6.15.0) and R (v4.3.1). Raw protein abundance quant values were log2 transformed and the distributions in each sample were both centred around 0 by median subtraction and width adjusted with quartile normalization. Samples were categorised into patient and control groups per brain region. Detected protein groups (n=4901 proteins) were filtered to ensure that at least two peptides were detected per protein, and each protein was detected in at least 70% of cases in at least one group (n=3790 proteins). The four groups were analysed using one-way analysis of variance (ANOVA) followed by Tukey’s post-hoc test, with multiple comparisons adjusted at a false discovery rate (FDR) threshold of *P* < 0.05.

Pathway enrichment analysis was performed using ENRICHR (accessed on 06/11/2025) with the Reactome database as the annotation source using Entrez gene IDs mapped to *Homo sapiens* [7, 25, 66]. Significantly increased and decreased proteins with a fold change greater than ±1.5 (0.58 log2 fold change) were compared against a custom background list containing all analysed proteins (n=3790) to account for dataset-specific detection bias. Enriched pathways with an adjusted *P*LJ<LJ0.05 were considered significant.

### Immunohistochemistry to identify inflammatory markers

To characterise inflammatory pathology and validate top altered pathways identified from the proteomics study, immunohistochemistry using FFPE sections was performed. All primary antibodies and antigen retrieval methods are provided in **Supplementary Table 5**. As previously described [48], 5µm-thick FFPE sections were deparaffinised and rehydrated through a series of Ethanol solutions to water. Antigen retrieval was performed using either 10mM Trisodium citrate (pH 6.0) or 10mM Tris 1mM EDTA (pH 9.0). All primary antibodies were diluted in Tris-buffered saline, 0.1% Tween 20® (TBST, pH 7.4) and were applied to the sections overnight at 4°C. A MACH 4 universal horse radish peroxidase (HRP) polymer probe kit (Cell Path) and 3,3’-diaminobenzidine (DAB, Biocare Medical) was used to visualise the primary antibodies. Sections were then counterstained with Mayer’s Haematoxylin and Scott’s tap water to blue nuclei.

### Quantification of inflammatory cell densities

Immunohistochemistry slides were scanned using a motorised Zeiss Axioscan 7 slide scanner and Zen imaging software. Slide scanned images were then imported into QuPath digital pathology image analysis software [4]. Slides which were not compatible for the Axioscan 7 slide scanner were viewed using an Olympus BX51 stereology microscope and StereoInvestigator software (MBF Bioscience, Williston, VT USA). The same cell density quantification protocol was applied using QuPath and StereoInvestigator. This involved outlining a contour of at least 10mm^2^ area, encompassing all cortical layers (I – VI) within BA17 and BA9, and counting DAB-positive cells within the contour. All cells immunoreactive for glial proteins, which were not localised within blood vessels, were counted. Since the mitochondrial TSPO antibody weakly labelled neuronal cells in all cases, only cells with a morphology typical of glial cells were counted. For patients with focal stroke-like lesions, cells immunoreactive for all markers were quantified in focal lesioned cortex and neighbouring non-lesioned cortex, with an average taken across the two areas. Cell density counts were independently verified by a second assessor.

### Multiplex immunofluorescence to identify mitochondrial proteins in microglia

To investigate changes to the expression of mitochondrial OXPHOS proteins within microglia in POLG-related epilepsy, a previously optimised multiplex immunofluorescence assay was adapted to identify the mitochondrial complex I subunit NDUFB8 (NADH:ubiquinone oxidoreductase subunit B8), complex IV subunit COXI (cytochrome *c* oxidase subunit I), and mitochondrial mass marker porin (voltage-dependent anion channel 1, VDAC1) within Iba-1-immunoreactive microglia (**Supplementary Table 6**) [15, 48]. A loss of NDUFB8 and COXI protein expression relative to porin have previously been reported in cortical inhibitory interneurons and astrocytes from patients with Alpers’ syndrome [15, 48, 49], indicative of impaired structure and function of complex I and complex IV [55, 67].

Briefly, this multiplex immunofluorescence assay involved antigen retrieval using 1mM EDTA (pH 8.0) in a pressure cooker for 40 minutes and blocking in 10% normal goat serum (NGS)-TBST for 1 hour at room temperature, followed by avidin and biotin block for 15 minutes each. The primary antibody cocktail (Iba-1, NDUFB8, COXI and Porin) was applied to the sections overnight at 4°C (**Supplementary Table 6**). The following day, a biotinylated anti-mouse IgG1 biotin antibody was applied to the sections for 30 minutes at room temperature to amplify the NDUFB8 primary antibody. Alexa Fluor-conjugated secondary antibodies were next applied for 2 hours at 4°C (**Supplementary Table 6**). To minimise autofluorescence, the sections were incubated with 70% ethanol for 1 minute, followed by 1% Sudan Black B solution (diluted in 70% Ethanol) for 10 minutes at room temperature. The sections were then mounted with ProLong^TM^ Gold antifade reagent and were dried at room temperature for 48 hours. FFPE tissues which were fixed in formalin for more than 1 year were not included in this immunofluorescence assay, since long formalin fixation durations are known to alter antigenicity [62]. All occipital cortex tissues were included in the same experimental batch, and all frontal cortex tissues were included in a second experimental batch.

### Three-dimensional analysis of microglia

Multiplex immunofluorescence sections were imaged using an inverted ZEISS LSM800 confocal microscope and ZEISS Zen (blue edition) software at x63 magnification (oil immersion lens), with a x3 electronic zoom, as previously described [49]. Z-stacking was performed which involved imaging microglia on the x-, y- and z-planes using a z-step size of 0.30µm, with two times line averaging [48]. At least 25 microglia were randomly imaged across the cortical layers within BA17 and BA9 per case, based on Iba-1 immunoreactive cell bodies and associated processes. Optimised image capture settings were maintained for all cases, per experiment.

All images were deconvolved using Huygens Essential software (version 24.04, Scientific Volume Imaging, The Netherlands, http://svi.nl), applying the same optimised deconvolution and microscopic parameters. Deconvolved images were then imported into Imaris (version 9.9.2) for three-dimensional analyses. Based on the intensity of the 405nm signal, Iba-1+ microglia were detected and were set as an ‘object’. The mean intensity of COXI (488nm), Porin (546nm) and NDUFB8 (647nm) signals were measured within the Iba1+ microglial ‘object’. The total volume (µm^3^) and surface area (µm^2^) of individual Iba-1+ microglia were also measured. Per cell, the mean intensity of NDUFB8 and COXI were log-transformed and divided by the log-transformed porin intensity, to normalise for mitochondrial mass. To determine the levels of NDUFB8 and COXI protein deficiencies within patient microglia, z-scores were calculated using the NDUFB8/Porin and COXI/Porin ratios, as previously described [44]. To infer changes to mitochondrial mass within patient microglia, z-scores were also calculated using the mean log-transformed porin intensity values.

### Statistical analysis

Neuropathology data were analysed using GraphPad Prism 10.0 (GraphPad Software, Inc., La Jolla, California) and R (v3.4.1 ). Normality was assessed using the Shapiro-Wilk test alongside visual inspection of Q-Q plots. Per brain region, cell densities were compared between patient and control groups using a Mann-Whitney U test. Data from the multiplex immunofluorescence OXPHOS assay were analysed using a linear mixed-effects model, with group included as a fixed effect and case as a random effect, accounting for the number of Iba-1LJ microglia analysed per case [36]. Statistical significance was set at α = 0.05.

## Results

### Clinical and demographic details

The patient cohort comprised of 12 patients with bi-allelic POLG-related disease (**Table 1**). In this cohort, the median age at disease onset was 10 years, with 6 patients presenting before 12 years of age. The disease duration ranged from 1 month to 16 years and the median age at death was 23.5 years (**Supplementary Table 1**). Of the 7 patients who died as young adults, 85.7% were female, whereas those dying in infancy or early childhood (n=5) showed a male predominance (80% male). Of the 7 different *POLG* variants identified in this cohort, p.[Ala467Thr] and p.[Trp748Ser] were the most common variants segregating with disease (**Table 1**). One or both of these variants were present in all individuals in this cohort.

Refractory status epilepticus was the main presenting symptom in all patients and was confirmed to have an occipital involvement in 11 individuals (**Table 1**). Cortical visual symptoms including blindness were reported in 8 patients, and 9 patients presented with stroke-like episodes.

### Occipital cortex shows an altered proteome in POLG-related epilepsy

To identify altered pathways that could contribute to the neuropathological basis of epilepsy in POLG-related disease, we compared the abundance of 3,790 proteins across 40 samples from patients with POLG-related epilepsy (n=9 occipital and n=10 frontal patient samples) and matched controls (n=10 occipital and n=11 frontal control samples) (**Fig. 1A, Supplementary Table 4**). Given that we were interested in identifying proteomic differences that were unique within the primary visual cortex in addition to those shared with the frontal cortex, both brain regions were compared between and within patient and control groups.

**Fig 1.**
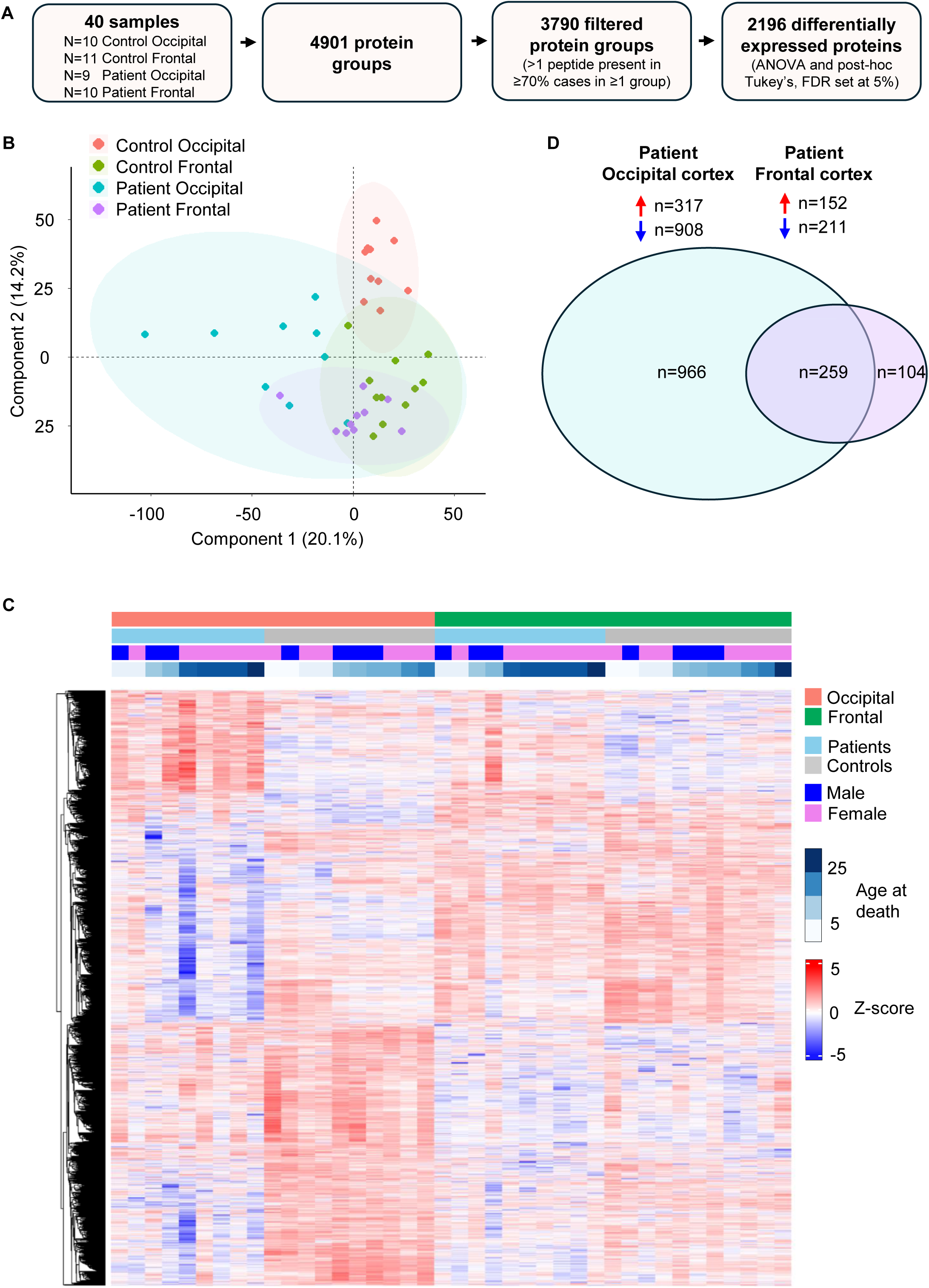
Quantitative mass spectrometry proteomics using occipital and frontal cortical tissues from patients with POLG-related epilepsy and matched control cases. **(A)** Experimental workflow and statistical analyses for LC-MS/MS is summarised. N=10 control occipital, n=11 control frontal, n=9 patient occipital and n=10 patient frontal cortex tissues were included (see Supplementary Table 1 – 3 for details). **(B)** Principal component analyses (PCA) demonstrate distinct differences between the patient and control groups, and between brain regions. **(C)** The Heatmap displays the 2,196 significantly differentially expressed proteins, with log2 fold changes normalised and expressed as z-scores. Rows were clustered using average-linked hierarchical clustering based on Euclidean distance to group proteins with similar expression profiles. The colour coded key provides details of patient and control cases, including sex and age at death. **(D)** Venn diagram summarises the total number of significantly differentially expressed proteins in patient tissues compared to control tissues, per brain region (Tukey’s test, 5% FDR, adjusted *P* < 0.05)

Principal component analysis showed distinct clustering of patients from the control cases in the occipital cortex, whereas separation between the patients and controls was less pronounced in the frontal cortex (**Fig. 1B**). Noticeable differences between occipital and frontal cortical samples of both patient and control groups were also observed due to brain region-specific proteomic differences.

ANOVA identified 2,196 proteins that differed significantly across the four groups (**Fig. 1C**). Post hoc multiple comparison testing identified 1,225 proteins with significantly altered abundance between patients with POLG-related epilepsy and controls in the occipital cortex, compared with 363 in the frontal cortex (**Fig. 1D**). Noticeably, a higher proportion of the total differentially expressed proteins in the patient occipital cortex group were decreased (74.12%) relative to proteins with an increased abundance (25.88%); a pattern that was less pronounced in the frontal cortex group. These data indicate more extensive proteomic alterations in the occipital cortex than in the frontal cortex in POLG-related epilepsy.

### Decreased abundance of mitochondrial OXPHOS and neuronal proteins in POLG-related epilepsy

Since the occipital cortex is particularly susceptible to seizures in POLG-related epilepsy [9], and severe neurodegeneration [13], we next included proteins that were significantly decreased (adjusted *P*<0.05 and Log2FC <-0.58) in patient occipital (n=535) and frontal (n=92) cortical regions for pathway analysis.

Analysis of the 20 most significantly decreased proteins in the occipital cortex of patients with POLG-related epilepsy (ranked by adjusted *P*-value) showed that 14 were core or accessory subunits of mitochondrial OXPHOS complex I (**Fig. 2A** and **Supplementary Table 7**). Of 540 mitochondrial proteins (defined by MitoCarta 3.0) detected and included in the analysis [43], 130 were differentially expressed in the patient occipital cortex, of which 86.15% were less abundant than in controls (**Supplementary Table 8**). In comparison, 44 mitochondrial proteins were differentially expressed in the frontal cortex.

**Fig. 2.**
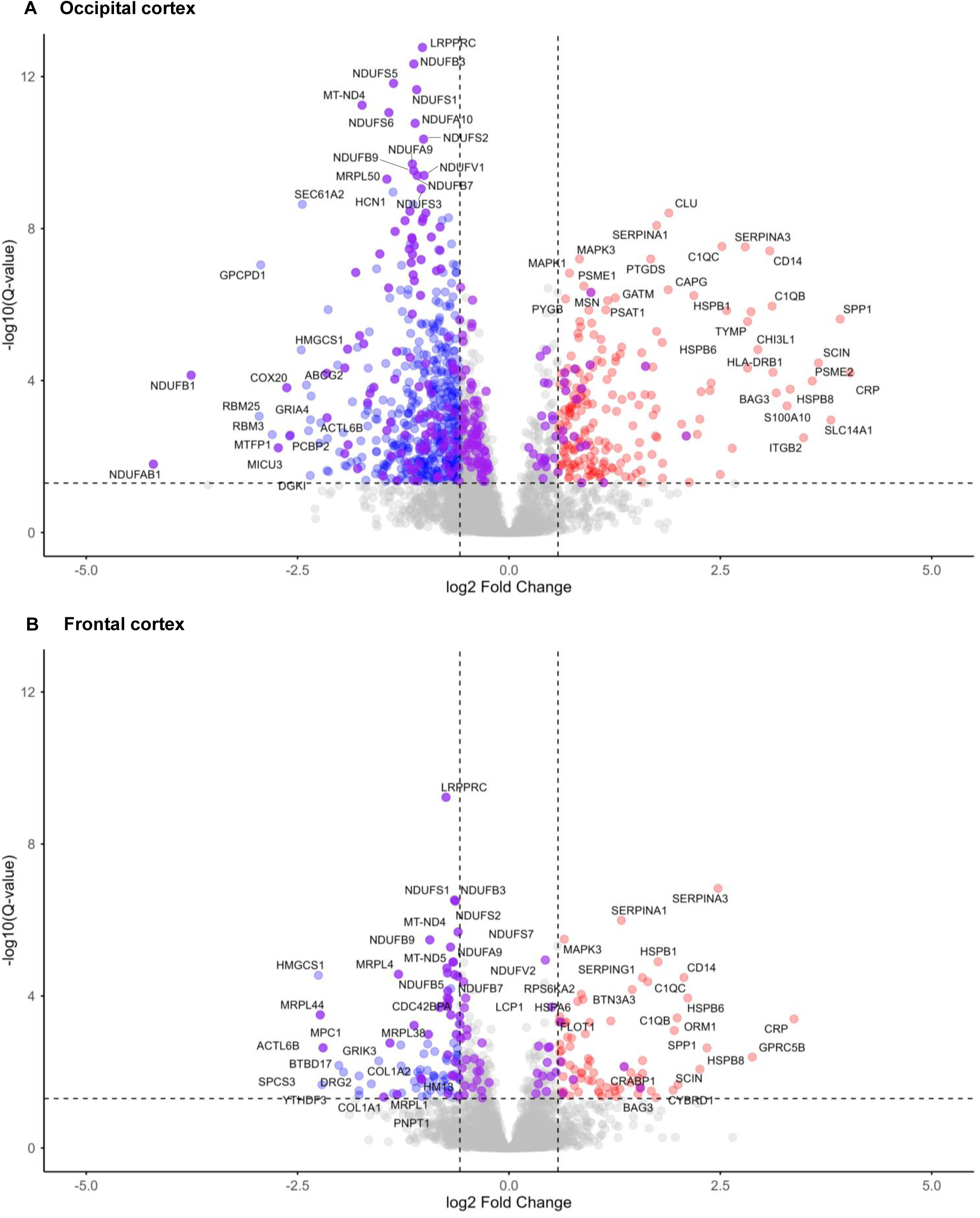
Differentially expressed proteins in the occipital and frontal cortex of patients with POLG-related epilepsy. Volcano plots demonstrate the proportion of significantly increased (upper right red circles) and significantly decreased (upper left blue circles) proteins with a log 2-fold change ± 0.58 (fold change ± 1.5) and -log10(Q value) >1.3, in POLG-related epilepsy tissues relative to matched controls. (**A**) n=201 increased and n=535 decreased proteins in patient occipital cortex tissues. (**B**) n=77 increased and n=92 decreased proteins in patient frontal cortex tissues. Significantly altered mitochondrial proteins are also highlighted in dark purple (MitoCarta 3.0)[43] for the occipital cortex (n=227) and frontal cortex (n=74). The gene IDs of the top differentially expressed proteins with the greatest increased and decreased log2 fold change are annotated, in addition to the top significantly altered proteins. Dashed vertical lines indicate the log2 fold change >0.58 and <-0.58, and dashed horizontal line indicates the significance threshold set at -log10 Q-value of 1.3 (i.e. adjusted *P* value < 0.05)

Pathway analyses revealed a total of 50 proteins linked to mitochondrial OXPHOS were significantly decreased in patient occipital cortex tissues, compared to 19 decreased in patient frontal cortex tissues (**Fig. 3, Supplementary Table 9**). Multiple subunits of complexes I, III and IV were decreased in the occipital cortex, whereas in the frontal cortex decreases were largely restricted to complex I, with only a single complex IV subunit affected. Proteins associated with mitochondrial translation pathways were less abundant in the occipital cortex, with smaller reductions in the frontal cortex (**Supplementary Tables 9–10**). These findings suggest more extensive OXPHOS impairment in the occipital cortex than in the frontal cortex in POLG-related epilepsy.

**Fig. 3.**
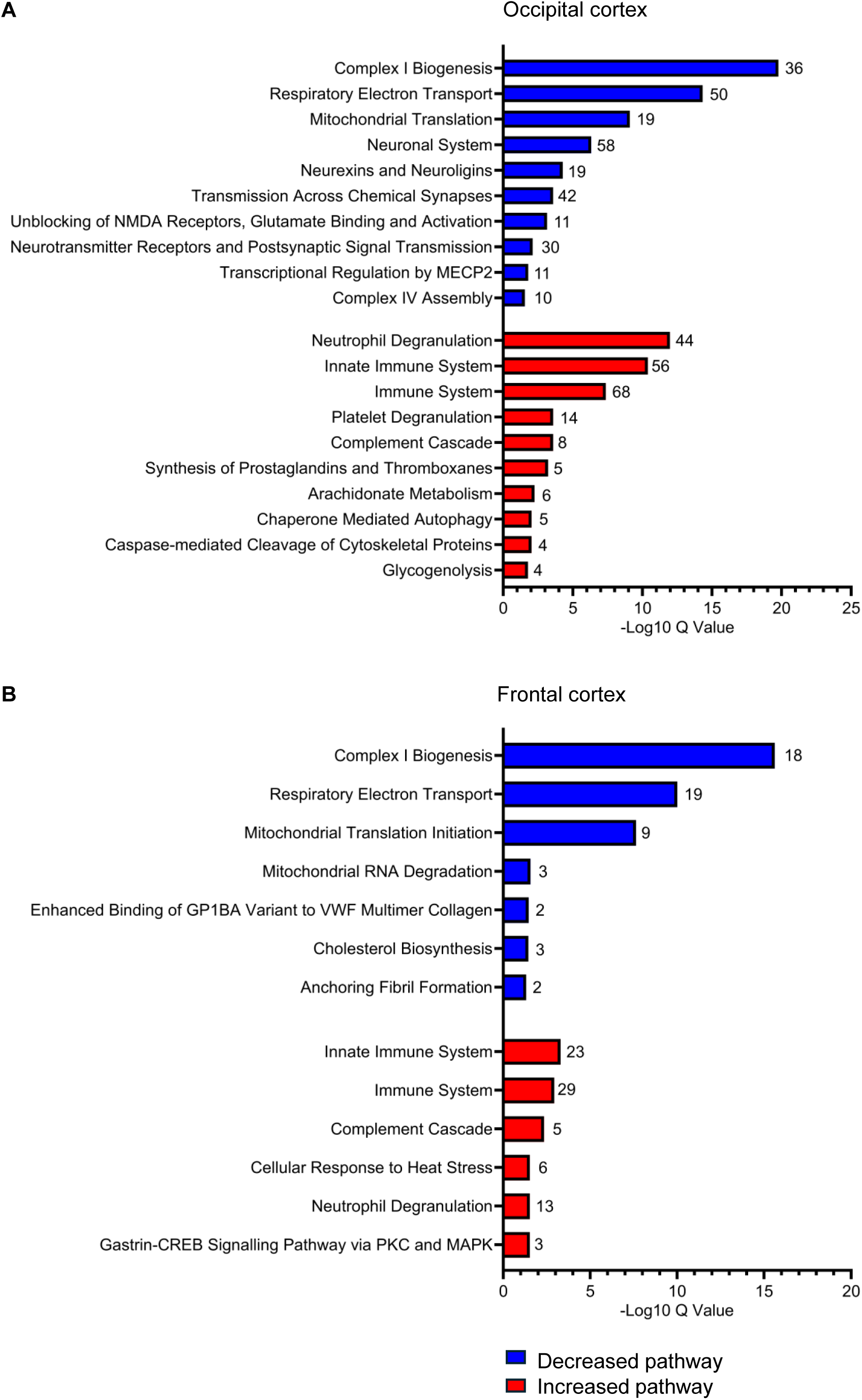
Reactome analysis identifies top altered pathways in the occipital and frontal cortex in POLG-related epilepsy. Pathway enrichment analyses were performed using ENRICHR and the Reactome database (accessed 06/11/2025). Analysis was conducted using significantly increased and decreased proteins (Q value < 0.05, and log2 fold change ± 0.58) against a custom background gene list composed of all analysed proteins (n=3790). The number of proteins used for pathway analyses included: n=535 decreased proteins and n=201 increased proteins in patient occipital cortex, and n=92 decreased proteins and n=77 increased proteins in patient frontal cortex. Top decreased pathways (blue bars) and top increased pathways (red bars) are shown, ranked by the greatest -10 log Q value for the (**A**) occipital cortex and (**B**) frontal cortex. Duplicate pathways are not displayed; see Supplementary Table 8 for all differentially expressed pathways. The total number of differentially expressed proteins associated with each pathway are displayed adjacent to the bars

Pathways uniquely downregulated in the occipital cortex of patients with POLG-related epilepsy included those related to neuronal and synaptic activity (**Fig. 3A**). This was accompanied by reduced abundance of inhibitory interneuron markers, including glutamate decarboxylase 1 and 2 (GAD65/67) and parvalbumin, as well as ionotropic glutamatergic receptor subunits and pre- and post-synaptic proteins, consistent with neuronal loss. Pathways associated with neuronal loss were not altered in the frontal cortex of patients with POLG-related epilepsy. In contrast, uniquely downregulated pathways in the frontal cortex were limited to cholesterol metabolism and haemostasis (**Fig. 3B; Supplementary Table 9**). In line with previous neuropathological reports [15, 49], these findings indicate greater degeneration of inhibitory interneurons and pyramidal neurons in the occipital cortex, supporting increased regional vulnerability in POLG-related disease.

### An innate immune signature is revealed in POLG-related epilepsy

We next examined the most upregulated proteins in POLG-related epilepsy, which showed increased abundance of innate immune proteins (**Fig.2A**). These included acute phase proteins (C-reactive protein, osteopontin and serpin A3), immune co-receptors (CD14 and HLA-DR), and components of the complement pathway, including C1QB. Heat shock proteins (HSPs), including HSPB8, HSPB6 and HSPB1, were also increased in both brain regions (**Supplementary Table 11**).

To investigate the functional relevance of upregulated proteins in POLG-related epilepsy, we performed pathway enrichment analysis using all proteins with increased abundance (adjusted *P* <0.05 and log2FC > 0.58) in each brain region. This identified enrichment of innate immune signalling pathways in the occipital cortex, including neutrophil and platelet degranulation, complement activation (**Fig. 3A**), and interleukin 12 and interferon-γ signalling (**Supplementary Table 9**).

Innate immune response proteins were also significantly enriched in patient frontal cortex tissues, including a shared upregulation of the complement cascade and neutrophil degranulation across both brain regions, albeit fewer proteins linked to these pathways were altered within the frontal cortex (**Fig. 3B** and **Supplementary Table 9**). These findings provide evidence of an innate immune signature shared across the occipital and frontal cortex in POLG-related epilepsy.

Patient occipital cortex tissues also demonstrated a significant increased abundance of proteins involved with glutathione conjugation (oxidative stress response), glycogen metabolism, and arachidonate metabolism, which plays a role in producing inflammatory mediators including prostaglandins [1, 2].

### High metabolic protein expression in control occipital cortex

To examine whether proteomic differences in the primary visual cortex contribute to its vulnerability to mitochondrial dysfunction in POLG-related disease, we analysed differentially expressed proteins between occipital and frontal cortex control samples.

Of the 843 proteins that were differentially expressed between the two brain regions, 589 proteins were more abundant in control occipital cortex (**Fig. 4A**). Pathway analysis of proteins with increased abundance (log2 fold change >0.58) showed enrichment of pathways related to mRNA processing, splicing and trafficking, indicating enhanced post-transcriptional regulation in the occipital cortex (**Supplementary Table 12**).

**Fig. 4.**
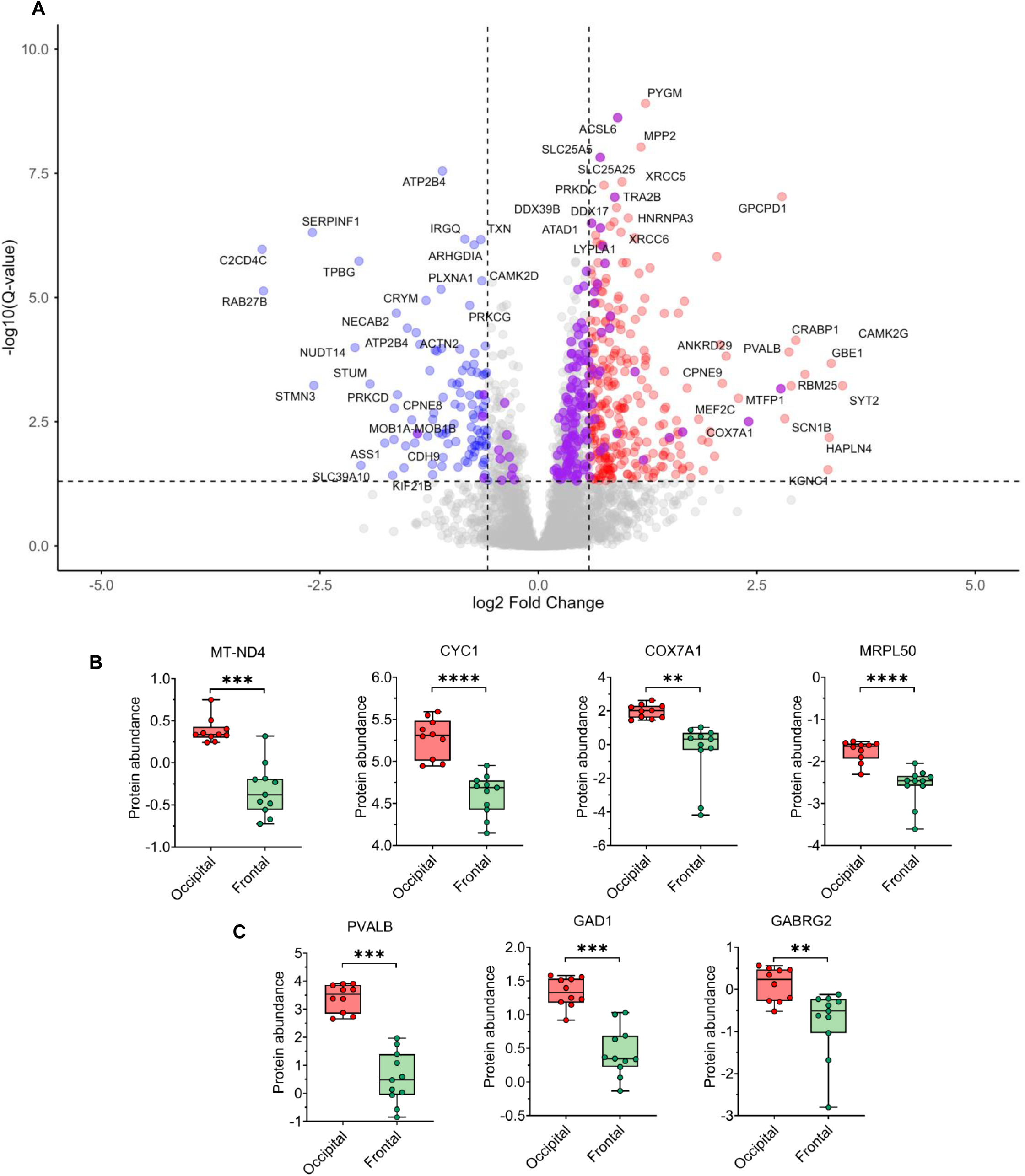
Proteomic differences in the occipital cortex versus frontal cortex of control cases. **(A)** Volcano plot demonstrates the proportion of significantly increased (upper right red circles, n=293) and significantly decreased (upper left blue circles, n=105) proteins in occipital cortex tissues compared to frontal cortex tissues of control cases (Q value < 0.05, and log2 fold change ± 0.58). Significantly altered mitochondrial proteins (n=147) are also highlighted in purple (MitoCarta 3.0).[43] The gene IDs of the top differentially altered proteins with the greatest increased and decreased log2 fold change are annotated, in addition to the top proteins with the highest -log10(Q-values). Dashed vertical lines indicate the log2 fold change >0.58 and <-0.58, and dashed horizontal line indicates the significance threshold set at -log10 Q-value of 1.3 (i.e. adjusted *P* value < 0.05). **(B)** Bar charts demonstrate selected mitochondrial OXPHOS protein subunits that have a significantly increased protein expression in control occipital versus frontal cortex, in addition to **(C)** inhibitory interneuron proteins. ** *P* < 0.01, *** *P* < 0.001, **** *P* < 0.0001

Utilising the MitoCarta 3.0 proteome to define mitochondrial proteins revealed 134 mitochondrial proteins were significantly upregulated in occipital versus frontal cortical tissues of the control group (**Supplementary Table 8**) [43]. This included 70 proteins directly associated with OXPHOS (**Fig. 4B**), of which 31 proteins had a log2 fold change >0.58. A significantly increased abundance of proteins involved in the synthesis of the inhibitory neurotransmitter gamma-aminobutyric acid (GABA) and parvalbumin (protein expressed by fast-spiking interneurons) was also identified (**Fig. 4C**). Together, these findings suggest that the increased metabolic demands of the primary visual cortex may be largely determined by the need for extensive inhibitory regulation. This may be an important contributor to the vulnerability of the primary visual cortex to metabolic dysfunction in primary mitochondrial disease and may partly explain the predilection of this brain region in POLG-related epilepsy.

### Severe microgliosis and astrogliosis in the occipital cortex in POLG-related epilepsy

Given the increased abundance of innate immune signalling proteins identified in POLG-related epilepsy, we next sought to validate these findings in FFPE cortical tissues (**Supplementary Table 4**). As neuroinflammation is predominantly mediated by microglia and astrocytes [26], we quantified the density of HLA-DR+ activated microglia and GFAP+ reactive astrocytes in patient and control tissues.

In line with previous reports [48], all patients showed increased GFAP+ and HLA-DR+ cell densities in the occipital cortex compared to controls (GFAP: *P* < 0.0001; HLA-DR: *P* < 0.05; **Fig. 5**). In contrast, GFAP+ and HLA-DR+ cell densities did not differ significantly between groups in the frontal cortex (*P* > 0.05), indicating less marked astrogliosis and microgliosis in this region. In both cortical regions, GFAP+ and HLA-DR+ cells delineated the borders of focal stroke-like lesions (**Fig. 5**).

**Fig. 5.**
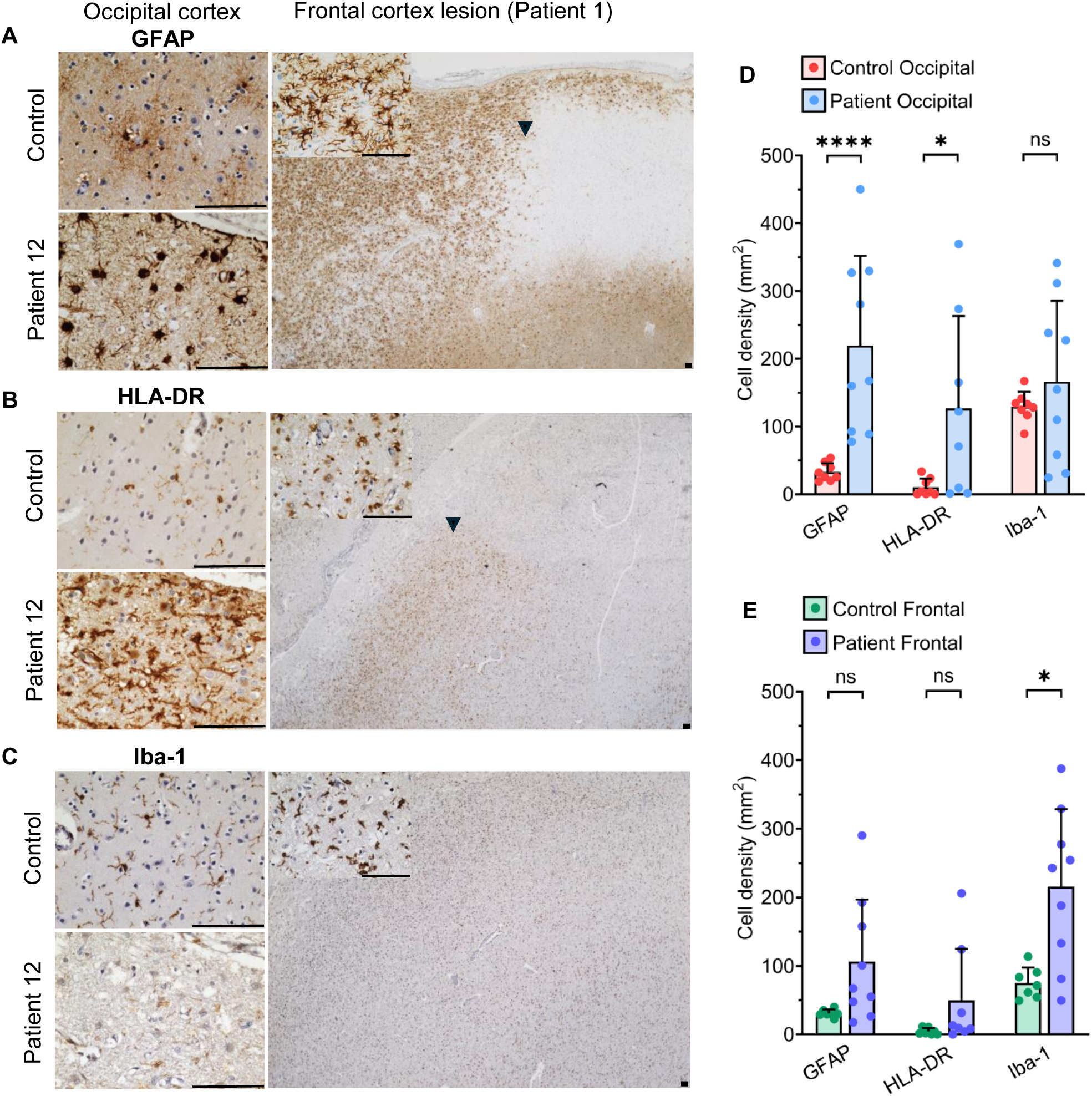
Reactive astrogliosis and microgliosis in POLG-related mitochondrial epilepsy. Representative immunohistochemistry images demonstrate the abundance of **(A)** GFAP-immunoreactive (+) astrocytes, **(B)** HLA-DR+ activated microglia, and **(C)** Iba-1+ microglia (pan-microglial marker) in control occipital cortex, POLG Patient 12 occipital cortex, and within a focal stroke-like lesion in the frontal cortex of Patient 1. Note the focal stroke-like lesion is demarcated by increased GFAP+ and HLA-DR+ immunoreactivity (black arrowhead highlights the border between lesioned and non-lesioned cortex) **(A – B)**, but the lesion is not clearly defined by Iba+ immunoreactivity **(C)**. Iba1+ immunoreactivity is also visibly decreased in the occipital cortex of Patient 12 **(C)**, despite high HLA-DR+ immunoreactivity **(B)**. All tissues were counterstained with Haematoxylin. Corresponding densities of cells immunoreactive for each protein are presented for the **(D)** occipital cortex and **(E)** frontal cortex. Circles represent individual cases. Data analysed using Mann-Whitney U test. * *P* < 0.05, ** P < 0.01, **** *P* < 0.0001, ns = non-significant (*P* > 0.05). Scale bars = 100 µm

To provide an insight to the proportion of microglia that are activated (i.e., HLA-DR+) in POLG-related epilepsy, we also quantified the number of cells immunoreactive for Iba-1, which is an ionised calcium-binding adaptor molecule expressed by microglia in addition to peripheral macrophages [8, 18]. Interestingly, this revealed a highly variable expression of Iba-1 in the POLG-related epilepsy patient group, with visibly diminished immunoreactivity of Iba-1 in patient cortical tissues affected by severe neurodegeneration (**Fig. 5C**), despite a high abundance of HLA-DR+ microglia (**Fig. 5B**). We also observed that focal stroke-like lesions were not clearly defined by Iba1+ immunoreactivity (**Fig. 5C**). These data suggest that Iba-1+ protein expression is impacted by POLG-related neurodegeneration and is not a reliable marker for characterising microgliosis in POLG-related epilepsy.

### Increased expression of inflammatory glial proteins in POLG-related epilepsy

To validate the most upregulated inflammatory proteins identified by proteomics, we quantified cells immunoreactive for acute phase proteins. Densities of C-reactive protein (CRP), serpin A3, osteopontin and CD14+ cells were increased in both the occipital and frontal cortex of patients (**Fig. 6**). Although labelling was predominantly glial, CRP, serpin A3 and osteopontin were also detected in neurons and were prominent within blood vessels (**Supplementary Fig. 1**). YKL40 (chitinase-3-like protein 1), an inflammatory glycoprotein expressed by activated astrocytes [5], was also increased in both cortical regions.

**Fig. 6.**
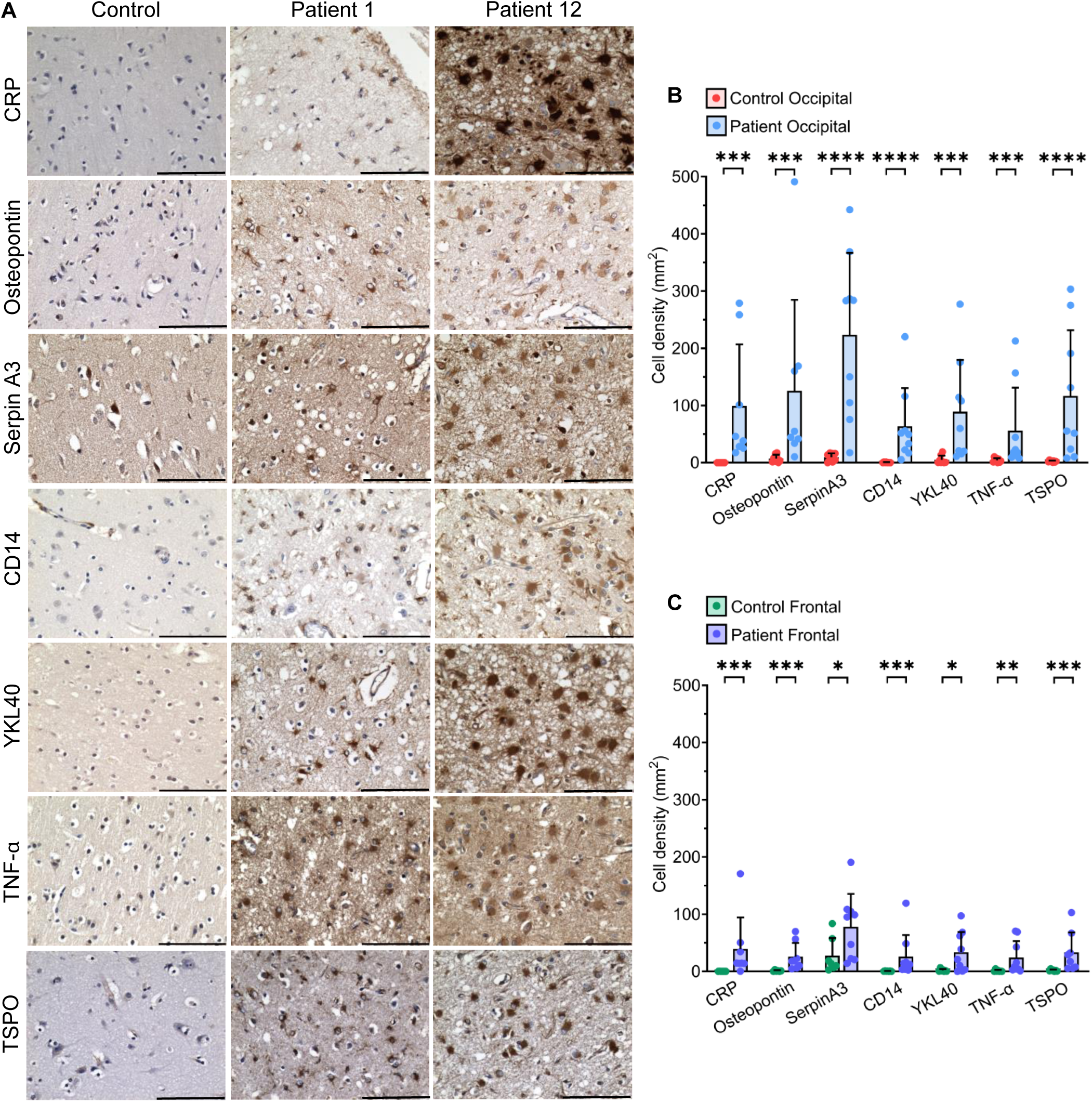
Increased expression of inflammatory glial proteins in POLG-related epilepsy. **(A)** Representative images demonstrate an increased abundance of C-reactive protein (CRP), osteopontin (acute phase glycophosphoprotein), serpin A3 (acute phase glycoprotein), CD14 (Toll-like receptor 4/TLR4 co-receptor), YKL40 (inflammatory glycoprotein), TNF-α (inflammatory cytokine) and TSPO (mitochondrial translocator protein) in occipital cortex patient tissues. Minimal expression of these inflammatory proteins was observed in control cases, except TSPO which weakly labelled neurons. Corresponding cell densities are provided for the **(B)** occipital cortex and **(C)** frontal cortex for patients and controls. Circles represent individual cases. Data analysed using Mann-Whitney U test. * *P* < 0.05, ** P < 0.01, *** *P* < 0.001, **** *P* < 0.0001. Scale bars = 100 µm

Although few inflammatory cytokines were detected in our proteomics study, likely due to the limited sensitivity of bulk LC-MS/MS for low abundant proteins, densities of tumour necrosis factor alpha (TNF-α)+ cells were increased in patient tissues (**Fig. 6**). TNF-α is an inflammatory cytokine that has been reported to promote neuronal hyperexcitability [40, 52].

To examine the relationship between mitochondrial dysfunction and glial activation, we quantified TSPO+ cells [47]. Densities of amoeboid TSPO-immunoreactive glial cells were increased in patient tissues, consistent with activation of both microglia and astrocytes (**Fig. 6**).

Since neutrophil degranulation was reported to be an enriched pathway in patient cortical tissues (**Fig. 3**), we quantified the density of cells immunoreactive for myeloperoxidase (MPO), an abundant protein stored within granules of neutrophils [45]. Analysis revealed a significantly increased density of MPO+ cells that appeared to be localised both within the brain parenchyma and associated vasculature in patient occipital cortex tissues (**Supplementary Fig. 2**). MPO+ cells had small circular cell bodies and did not appear to be glial or neuronal cells based on morphology.

Collectively, these data provide evidence of an increased density of cells expressing pro-inflammatory markers in occipital and frontal cortical tissues from patients with POLG-related epilepsy.

### Microglial oxidative phosphorylation protein deficiencies

To investigate whether POLG patient microglia demonstrate altered expression of mitochondrial OXPHOS proteins, indicative of complex I and complex IV OXPHOS dysfunction, we adapted a multiplex immunofluorescence assay to quantify NDUFB8 and COXI expression within Iba-1+ microglia, using porin as a mitochondrial mass marker [15, 48].

In the occipital cortex, NDUFB8 and COXI intensities (normalised to porin) were decreased in patient microglia compared with controls (*P* < 0.001; **Fig. 7**). Iba-1+ microglia in the frontal cortex showed similar decreases in NDUFB8 and COXI abundance. These findings indicate impaired OXPHOS function in microglia in both the occipital and frontal cortices in POLG-related epilepsy. To assess mitochondrial mass in microglia, porin intensity was compared between patients and controls. No significant differences were observed in either cortical region (*P* > 0.05), although increased porin intensity was evident in a subset of patients (**Fig. 7**).

**Fig. 7.**
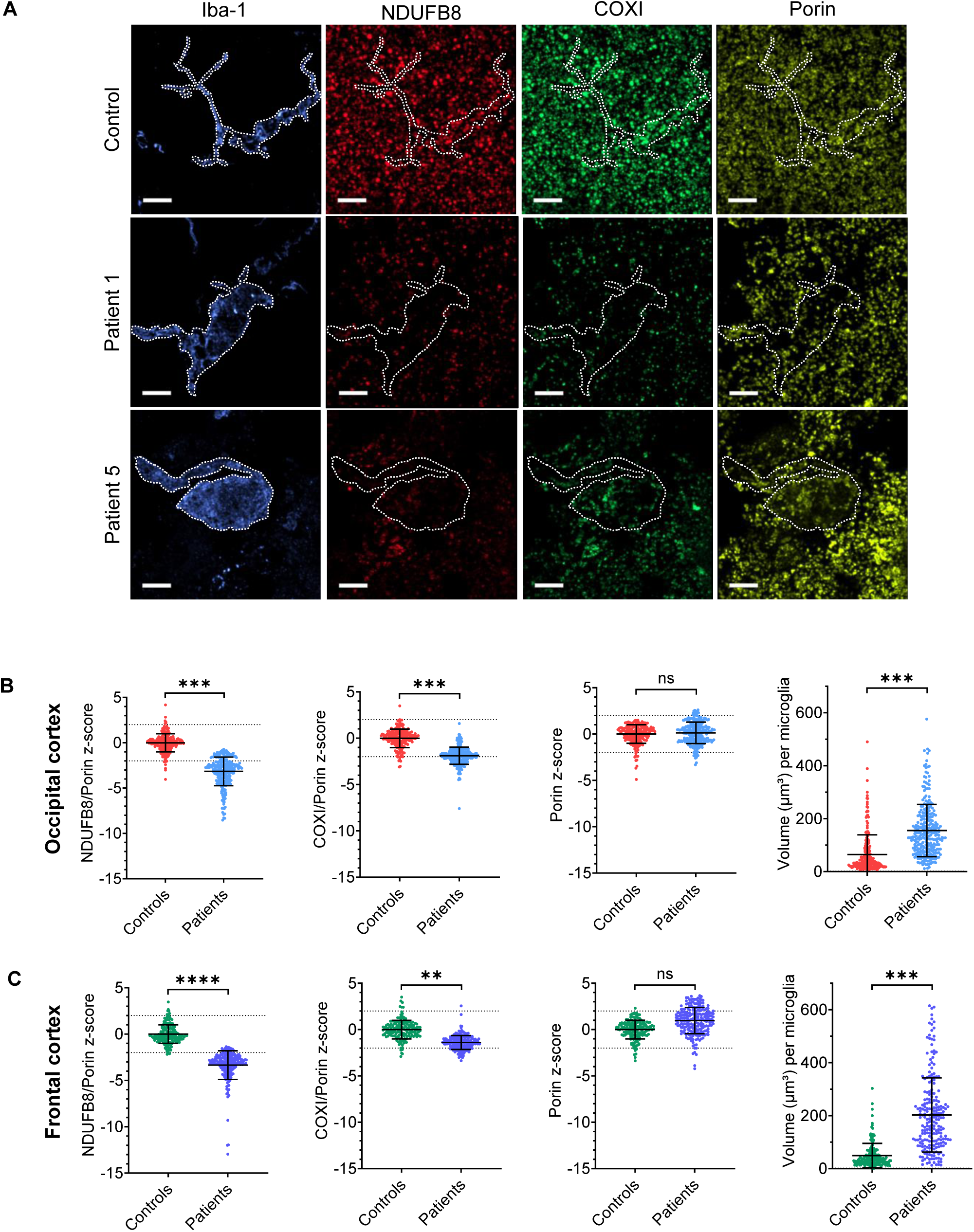
Mitochondrial oxidative phosphorylation protein deficiencies in POLG-related epilepsy patient microglia. **(A)** Representative images demonstrate a decreased intensity of NDUFB8 (complex I subunit) and COXI (complex IV subunit) within z-stacked POLG patient Iba-1+ microglia, relative to the intensity of Porin (mitochondrial mass marker). Scale bars = 10 µm. Graphs demonstrate z-scores for the intensity of NDUFB8 and COXI, normalised to Porin, for control and patient microglia in the **(B)** occipital cortex and **(C)** frontal cortex, in addition to the total volume of individual microglia (µm^3^). Circles represent individual microglia from n=8 controls and n=8 patients in the occipital cortex, and n=7 controls and n=8 patients in the frontal cortex. Data analysed using linear mixed effects model adjusted for multiple comparisons, with group included as a fixed effect and case as a random effect. **** *P* < 0.0001, *** *P* < 0.001, ** *P* < 0.01, ns = non-significant (*P* > 0.05)

Microglia are known to undergo morphological changes upon activation [35, 53]. Thus, we next quantified the surface area and volume of individual three-dimensional patient and control Iba-1+ microglia as an indicator of microglial activation. Both surface area and volume were increased in patient microglia compared with controls in the occipital and frontal cortices (*P* < 0.001; **Fig. 7**). Patient microglia also exhibited enlarged cell bodies with thickened, retracted processes, in contrast to the smaller, ramified morphology of control microglia. Taken together, these findings indicate diffuse microglial activation is coupled with reduced abundance of mitochondrial OXPHOS proteins in the cortex in POLG-related epilepsy.

## Discussion

The neuropathological mechanisms underlying POLG-related epilepsy remain poorly defined. Elucidating these mechanisms is essential for the development of effective therapies for this refractory condition. Here, we analysed post-mortem cortical tissue using complementary proteomic and immunohistochemical approaches. We identify an immunometabolic signature characterised by decreased abundance of mitochondrial proteins alongside increased innate immune and inflammatory proteins, consistent with neuroinflammation. This immunometabolic signature was observed in both the occipital and frontal cortex, despite the frontal cortex being comparatively preserved, suggesting that these proteomic alterations are not solely attributable to end-stage neurodegeneration. These findings suggest that mtDNA depletion-driven OXPHOS dysfunction may initiate both seizure activity and an innate neuroinflammatory process. Seizure activity likely further depletes ATP and disrupts the ionic mitochondrial membrane potential, exacerbating metabolic stress and promoting neuroinflammation. We also demonstrate enrichment of mitochondrial OXPHOS proteins and inhibitory interneurons in the control primary visual cortex relative to the frontal cortex, which may contribute to the selective vulnerability of this region in POLG-related disease.

Discovery mass spectrometry has previously been applied to analyse the proteome of post-mortem brain tissues from patients with epilepsy and POLG-related mitochondrial recessive ataxia syndrome (MIRAS) [20, 38], but this study is the first to focus on brain region-specific proteomic alterations in POLG-related mitochondrial epilepsy. The inclusion of tissue from two brain regions that are differentially affected by seizures and neurodegeneration enabled the identification of both shared and distinct proteomic alterations in the occipital and frontal cortex that we speculate could be associated with different stages of the disease process. The pathological changes observed in the occipital cortex likely reflect advanced seizure-associated neurodegeneration, whereas those in the frontal cortex may represent an earlier disease stage preceding end-stage neurodegeneration.

We identified a marked decrease in mitochondrial protein abundance in patient occipital cortex samples, with less pronounced changes in the frontal cortex. These data indicate more severe OXPHOS deficiency and mitochondrial disease pathology in the occipital cortex in POLG-related epilepsy. It is possible that this underlies greater neuronal and glial dysfunction, contributing to the establishment of aberrant, hyperexcitable cortical network activity within the primary visual cortex. Although we acknowledge that differences in the cellular composition of the occipital cortex in POLG-related disease could contribute to reductions in OXPHOS components, due to lower interneuron density and increased abundance of glycolytic glial cells, we have previously demonstrated significant OXPHOS protein deficiencies in the remaining interneurons and pyramidal neurons of the occipital cortex in POLG-related epilepsy, with deficiencies of greater magnitude than those observed in the frontal cortex [15, 48, 49]. Furthermore, we presently report OXPHOS protein deficiencies within patient Iba-1+ microglia. Taken together, these findings support more extensive OXPHOS impairment in the occipital cortex than the frontal cortex in POLG-related epilepsy, although the magnitude of this difference may be partly influenced by greater neuronal loss in the occipital cortex.

We also identified marked inflammatory changes across occipital and frontal cortices. These included an upregulation of innate immune pathways including the complement system, and increased abundance of acute phase proteins such as C-reactive protein, which may represent a potential biomarker of POLG-related epilepsy. Noticeably, neutrophil degranulation was also enriched, supported by increased density of MPO+ cells in patient tissues. MPO is abundant in neutrophil granules and contributes to ROS production and inflammatory signalling [12, 45]. However, proteins associated with neutrophil degranulation are also expressed by microglia [54]. Therefore, enrichment of this pathway may, in part, reflect increased microglial activation.

Although our bulk proteomics study was not sensitive enough to detect cytokines, we identified increased abundance of TNF-α in patient tissues. TNF-α can modulate neuronal receptor expression, decreasing GABAergic signalling while enhancing AMPA receptor expression, thereby promoting hyperexcitability [40, 52]. Taken together, these findings raise the possibility that inflammatory signalling could contribute to seizure activity in POLG-related epilepsy suggesting that an inflammatory component to the seizure phenotype may represent a potential therapeutic target. However, this warrants further investigation using suitable *in vitro* and *in vivo* models of POLG disease pathology.

Inflammation in POLG-related epilepsy may arise secondarily to neural dysfunction, neurodegeneration and seizure activity, which could explain why neuroinflammatory pathology was more pronounced in the occipital cortex than in the frontal cortex in POLG-related epilepsy patients. However, the presence of inflammatory changes in the frontal cortex suggests that inflammation is not solely a consequence of neurodegeneration but may also be driven by the underlying metabolic dysfunction in neurons and glia. Supporting this, pathogenic *POLG* variants have been linked to aberrant anti-viral response in patient cells [20], and a *Polg* mouse model demonstrates an elevated type I interferon response [31, 59].

We observed increased abundance of hypertrophic, activated microglia with reduced mitochondrial OXPHOS protein levels. As microglia undergo a metabolic switch from OXPHOS to glycolysis upon activation [11, 37], impaired OXPHOS may contribute to microglial activation and promote a pro-inflammatory phenotype. This may account for the presence of inflammatory pathology in the frontal cortex, where neurodegeneration is less pronounced. However, the precise impact of pathogenic *POLG* variants on microglial function remains unknown. We also observed reduced Iba-1 protein abundance in patient microglia within areas affected by severe cortical neurodegeneration. Similar reductions of Iba-1 have been reported in association with hepatic dysfunction [32], which is a common manifestation of early-onset POLG-related disease [24], in addition to age-associated subcortical white matter lesions [61].

The selective vulnerability of the primary visual cortex to seizures in POLG-related epilepsy remains poorly understood [9]. The high density of metabolically demanding parvalbumin-expressing interneurons in this region has been proposed to confer increased susceptibility to mitochondrial dysfunction [21, 22, 49]. Given their critical role in regulating neuronal network activity [46, 51], degeneration of these interneurons may contribute to the occipital focus of POLG-related epilepsy [49]. Consistent with this, we identified enrichment of metabolic proteins, particularly those involved in mitochondrial OXPHOS, in the primary visual cortex of control tissues. This was accompanied by an increased abundance of proteins involved in mRNA processing and translation, as well as GABAergic interneuron markers, including parvalbumin. Thus, the greater abundance of mitochondrial proteins in the occipital cortex may, in part, reflect the higher density of parvalbumin-expressing interneurons in this region. Other supportive evidence for a high metabolic demand in the occipital cortex comes from term infants exposed to profound hypoglycaemia, where differential injury to the visual cortex associated with epilepsy is observed [56]. Together, these findings suggest that the visual cortex has a high metabolic demand, such that depletion of mitochondrial proteins may have a disproportionate impact on neuronal and glial cell function.

Our proteomics dataset provides an invaluable resource for determining whether both *in vitro* and *in vivo* models of POLG-related pathology recapitulate the immunometabolic signature observed in brain tissues obtained from patients with POLG-related epilepsy. However, our findings have some caveats, including that they reflect end-stage disease pathology. The bulk tissue proteomics study was also limited by the inability to distinguish cell type-specific changes and the findings may therefore be confounded by alterations in cell type composition, which could be resolved using spatial proteomics. However, we have provided evidence of mitochondrial OXPHOS protein deficiencies within specific cell types [48–50], and have validated the top upregulated immune proteins using immunohistochemistry, confirming increased protein abundance within individual glial cells. While the proteomic coverage in this study may have limited the detection of low-abundance proteins, including some important mitochondrial proteins, the total number of identified proteins is comparable with published untargeted LC-MS/MS studies of post-mortem human brain tissue [19, 39, 60]. Furthermore, although post-mortem intervals (PMI) may have affected proteomic coverage in both patient and control tissues, the proteomics cohort was matched as closely as possible for PMI through the inclusion of suitable control tissues obtained from multiple brain banks. This represents an unavoidable limitation of studies using post-mortem tissues from young individuals. Moreover, although covariates including age at death, disease duration and sex were not included in our analysis, due to limited sample size, grouped level analyses has allowed for the identification of proteomic and neuropathological alterations that are common across the spectrum of early-onset POLG-related disease. Future studies comparing these findings with other mitochondrial diseases, including late-onset POLG-related disease, would be important for establishing whether the pathologies observed are disease-specific or have a broader relevance for other epilepsies where mitochondrial dysfunction is implicated.

In summary, this study provides insight into the metabolic and inflammatory mechanisms underlying POLG-related epilepsy. Further investigation of immune pathways in disease-relevant neuronal-glial models will help determine whether the observed inflammatory changes are causal or secondary.

## Data availability

The data supporting the findings of this study are available from the corresponding authors upon reasonable request.

## Supplementary material

Supplementary material is available.

## Declarations

### Compliance with ethical standards

Ethical approval was provided by Newcastle University Ethics Committee (Ref: 32755/2023) and the individual brain banks, including Newcastle Brain Tissue Resource (NBTR) 19/NE/0008, Brain UK (23/014), Oxford Brain Bank (15/SC/0639), East of Scotland Research Ethics Service REC1, NeuroBioBank (NBB) and Norwegian Regional Ethics Committee (2019/632). The UK Brain banks are UK Human Tissue Authority-approved and informed consent was obtained.

### Supplementary information

Supplementary material is available.

## Acknowledgements

We thank all patients and families for the donation of tissue samples for research. Tissue for this study was provided by the Newcastle Brain Tissue Resource which is funded in part by a grant from the UK Medical Research Council (G0400074), by NIHR Newcastle Biomedical Research Centre and as part of the Brains for Dementia Research Programme jointly funded by Alzheimer’s Research UK and Alzheimer’s Society. Tissue samples were also obtained from: the Edinburgh Brain and Tissue Bank which is funded by the Medical Research Council; the UK Brain Archive Information Network (BRAIN UK) (centres based in Southampton and Oxford) which is supported by Brain Tumour Research and has been established with the support of the British Neuropathological Society and the Medical Research Council; the Neuro-SysMed Center Brain Bank, Haukeland University Hospital, Bergen, Norway, and the National Institutes of Health NeuroBioBank at the University of Maryland, Baltimore. We thank the NHS Highly Specialised Mitochondrial laboratory team at Newcastle upon Tyne Hospitals NHS Foundation Trust for providing molecular diagnoses for patients included in this study. We are grateful to the staff of the Bioimaging Facility, Faculty of Medical Sciences, Newcastle University, for their help with confocal microscopy, image analyse advice and slide scanner service. We also thank the staff of the Newcastle University Protein and Proteome Analysis Facility, Faculty of Medical Sciences, Newcastle University, for performing the untargeted proteomics.

## Author contributions

L.A.S., D.E. and R.M. designed the study. L.A.S., M.W., E.M.E., Z.J., L.A., C.H., J.D. and M.A. conducted neuropathology experiments and performed data analyses. A.L.S. cut the FFPE tissue blocks and prepared sections for the study. R.W.T., O.H. and C.T. provided post-mortem tissue, molecular genetic and clinical data. P.P performed mass spectrometry. L.A.S., G.H. and P.P. analysed the proteomic data. L.A.S. wrote the first draft of the manuscript. All authors contributed to the interpretation of the data, critically revised the manuscript and gave their final approval.

