## Supplementary Information for "Neuroinflammation and metabolic dysfunction in POLG-related mitochondrial epilepsy"

**Supplementary Table 1** Demographic details for the POLG-related epilepsy post-mortem tissue cohort

| Case | ID | Brain Bank | Age at death | Age of disease onset | Disease duration | Sex | Pathogenic bi-allelic *POLG* variant | | Cause of death | Brain weight | Post-mortem interval | Formalin fixation duration |
|---|---|---|---|---|---|---|---|---|---|---|---|---|
| | | | | | | | Protein | DNA | | | | |
| Pt.01 | BBN003.36915 | NBTR | 3 y | 2 y | 9 m | M | p.[Cys418Arg]/p.[Ala467Thr] | c.1252T>C /c.1399G>A | Multi organ failure | 1106 g | 18 h | 1 m |
| Pt.02 | BBN004.33624 | The Oxford Brain Bank | 3 y | 3 y | 1 m | F | p.[Ala467Thr]/p.[Arg852Cys] | c.1399G>A/c.2554C>T | Complication of disorder | 1159 g | 27 h | 1 m |
| Pt.03 | 5317 | NBB | 11 y | 2 y | 9 y | M | p.[Ala467Thr]/p.[Gly848Ser] | c.1399G>A/c.2542G>A | Complication of disorder | N/A | 21 h | X |
| Pt.04 | 6397 | NBB | 13 y | 1 y | 12 y | M | p.[Leu424fs] / p.[Ala467Thr] | c.1270_1271 delCT/c.1399G>A | Complication of disorder | 437 g | 24 h | 4 y |
| Pt.05 | WS9A | Bergen | 13 y | 2 y | 11 y | M | p.[Trp748Ser]/p.[Trp748Ser] | c.2243G>C/c.2243G>C | Status epilepticus | | | |
| Pt.06 | BBN003.33660 | NBTR | 23 y | 18 y | 5 y | F | p.[Ala467Thr]/p.[Ala467Thr] | c.1399G>A/c.1399G>A | Status epilepticus | 1263 g | 32 h | 1 m |
| Pt.07 | 2001.1701 | NBTR | 24 y | 20 y | 4 y | F | p.[Ala467Thr]/p.[Trp748Ser] | c.1399G>A/c.2243G>C | Suppurative tracheobronchitis | N/A | 83 h | 4.5 m |
| Pt.08 | BBN003.37949 | NBTR | 24 y | 8 y | 16 y | F | p.[Ala467Thr]/p.[Arg597Trp] | c.1399G>A/c.1789c>t | Intractable seizures | 1220 g | 75 h | X |
| Pt.09 | WS2A | Bergen | 24 y | 15 y | 9 y | F | p.[Trp748Ser]/p.[Trp748Ser] | c.2243G>C/c.2243G>C | Fulminant liver failure (not valproate induced) | N/A | N/A | X |
| Pt.10 | WS10A | Bergen | 24 y | 16 y | 8 y | F | p.[Trp748Ser]/p.[Trp748Ser] | c.2243G>C/c.2243G>C | Status epilepticus | N/A | 87 h | N/A |
| Pt.11 | WS8A | Bergen | 28 y | 12 y | 16 y | M | p.[Trp748Ser]/p.[Trp748Ser] | c.2243G>C/c.2243G>C | Status epilepticus | | | |
| Pt.12 | BBN003.32572 | NBTR | 28 y | 16 y | 12 y | F | p.[Ala467Thr]/p.[Trp748Ser] | c.1399G>A/c.1399G>A | Status epilepticus | 1352 g | 64 h | 1m |

**Abbreviations**: *NBTR* Newcastle Brain Tissue Resource; *F* Female; *M* Male; *y* years; *h* hours; *m* months; *g* grams.
Bergen Brain Bank: Neuro-SysMed Center Brain Bank, Haukeland University Hospital, Bergen, Norway.
*X* FFPE tissue was unavailable for the neuropathology study. *N/A* Not available

**Supplementary Table 2** Control post-mortem tissue cohort for the proteomics study

| Case ID | | Brain Bank | Age at death | Sex | Cause of death | Post-mortem interval | Previous publications |
|---|---|---|---|---|---|---|---|
| Ct.01 | 4455 | NBB | 1 y | F | Drowning | 39 h | |
| Ct.02 | 5891 | NBB | 1 y | M | Sudden infant death syndrome | 24 h | |
| Ct.05 | 4394 | NBB | 3 y | F | Fever, undetermined | 12 h | |
| Ct.06 | 4441 | NBB | 3 y | F | Acute and chronic asthma | 15 h | |
| Ct.08 | 5334 | NBB | 12 y | M | Hanging, suicide | 15 h | 1,2,3 |
| Ct.09 | 5376 | NBB | 12 y | M | Hanging, suicide | 19 h | |
| Ct.10 | 1024 | NBB | 13 y | M | Cardiac arrhythmia | 16 h | |
| Ct.11 | 5309 | NBB | 14y | F | Streptococcal toxic shock syndrome | 8 h | 1,2,3 |
| Ct.14 | BBN003.29560 | NBTR | 18 y | F | MDMA toxicity, cardiac arrest | 81 h | 1,2,3,4 |
| Ct.16 | 5646 | NBB | 20 y | F | Reactive airway disease | 23 h | |
| Ct.18 | BBN004.35803 | The Oxford Brain Bank | 28 y | F | Suicide | 104 h | |

**Abbreviations**: *NBB* NeuroBioBank; *NBTR* Newcastle Brain Tissue Resource; *F* Female; *M* Male; *y* years; *h* hours; *d* days.
*X* FFPE tissue was unavailable for the neuropathology study. *N/A* Not available.
Note: * Only frontal cortical tissue from Ct.18 was included in the proteomics study (primary visual cortical tissue was unavailable).

**Supplementary Table 3** Control post-mortem tissue cohort for the neuropathology study

| Case ID | | Brain Bank | Age at death | Sex | Cause of death | Formalin fixation duration (FFPE) | Post-mortem interval | Previous publications |
|---|---|---|---|---|---|---|---|---|
| Ct.03 | N272P.19 | BRAIN UK (UHS) | 2 y | M | Sudden infant death syndrome | 3 d | 144 h | |
| Ct.04 | N.71A.18 | BRAIN UK (UHS) | 3 y | M | Multiple injuries. | 7 d | 144 h | |
| Ct.07 | N94F.17 | BRAIN UK (UHS) | 6 y | F | Aortitis; hemopericardium | 5 d | 168 h | |
| Ct.12 | BBN.2448 | EBTR | 16 y | F | Sudden cardiac death | 6 d | 49 h | 1,2 |
| Ct.13 | BBN.2338 | EBTR | 16 y | M | Suspension by ligature | 4 d | 47 h | 1,2,3,4 |
| Ct.15 | BBN001.26976 | EBTR | 19 y | M | Unascertained, sudden death | N/A | 101 h | |
| Ct.17 | BBN.2442 | EBTR | 24 y | F | Suspension by ligature | 9 d | 47 h | 1,2,3,4 |
| Ct.18* | BBN004.35803 | The Oxford Brain Bank | 28 y | F | Suicide | N/A | 104 h | |

**Abbreviations**: *UHS* University Hospital Southampton; *NBTR* Newcastle Brain Tissue Resource*; EBTR* Edinburgh Brain Tissue Resource; *F* Female; *M* Male; *y* years; *h* hours; *d* days. *X* FFPE tissue was unavailable for the neuropathology study. *N/A* Not available.

**Supplementary Table 4** Summary of tissue included in the neuropathology and proteomics studies

| Case ID | | Brain Region | Neuropathology study (FFPE) | Proteomics study (Frozen) |
|---|---|---|---|---|
| Pt.01 | BBN003.36915 | Occipital, Frontal | + | + |
| Pt.02 | BBN004.33624 | Occipital, Frontal | + | + |
| Pt.03 | 5317 | Occipital, Frontal | – | + |
| Pt.04 | 6397 | Occipital, Frontal | + | + |
| Pt.05 | WS9A | Occipital, Frontal | + | – |
| Pt.06 | BBN003.33660 | Occipital, Frontal | + | + |
| Pt.07 | 2001.1701 | Occipital, Frontal | + | + |
| Pt.08 | BBN003.37949 | Occipital, Frontal | – | + |
| Pt.09 | WS2A | Frontal | – | + |
| Pt.10 | WS10A | Occipital, Frontal | + | + |
| Pt.11 | WS8A | Occipital, Frontal | + | – |
| Pt.12 | BBN003.32572 | Occipital, Frontal | + | + |
| Ct.01 | 4455 | Occipital, Frontal | – | + |
| Ct.02 | 5891 | Occipital, Frontal | – | + |
| Ct.03 | N272P.19 | Occipital, Frontal | + | – |
| Ct.04 | N.71A.18 | Occipital, Frontal | + | – |
| Ct.05 | 4394 | Occipital, Frontal | – | + |
| Ct.06 | 4441 | Occipital, Frontal | – | + |
| Ct.07 | N94F.17 | Occipital, Frontal | + | – |
| Ct.08 | 5334 | Occipital, Frontal | – | + |
| Ct.09 | 5376 | Occipital, Frontal | – | + |
| Ct.10 | 1024 | Occipital, Frontal | – | + |
| Ct.11 | 5309 | Occipital, Frontal | – | + |
| Ct.12 | BBN.2448 | Occipital, Frontal | + | – |
| Ct.13 | BBN.2338 | Occipital, Frontal | + | – |
| Ct.14 | BBN003.29560 | Occipital, Frontal | – | + |
| Ct.15 | BBN001.26976 | Occipital, Frontal | + | – |
| Ct.16 | 5646 | Occipital, Frontal | – | + |
| Ct.17 | BBN.2442 | Occipital, Frontal | + | – |
| Ct.18 | BBN004.35803 | Occipital, Frontal | + | + (Frontal only) |

Neuropathology study: n=10 POLG patients, n=8 control cases. Proteomics study: n=10 POLG patients, n=11 control cases.

**Supplementary Table 5** Immunohistochemistry primary antibodies

| Primary antibody | Target | Host (isotype) | Chromogen dilution | Antigen retrieval | Antibody supplier | Catalogue number (RRID) |
|---|---|---|---|---|---|---|
| Glial fibrillary acidic protein (**GFAP**) | Reactive astrocytes | Rabbit (IgG) | 1:15,000 | | DAKO | Z0334 (RRID:AB_10013382) |
| Human leukocyte antigen-DP, DQ, DR (**HLA-DR**) | Antigen-presenting cell surface glycoprotein receptor; increased expression by activated microglia and monocytes | Mouse (IgG1) | 1:1000 | | DAKO | M0775 (RRID:AB_2313661) |
| Chitinase 3-like 1 protein (**YKL40**) | Inflammatory glycoprotein; increased expression in reactive astrocytes | Rabbit (IgG) | 1:1000 | | Abcam | Ab77528 (RRID:AB_2040911) |
| Cluster of differentiation 14 (**CD14**) | Toll-like receptor 4 (TLR4) co-receptor; increased expression by activated microglia, monocytes and some reactive astrocytes | Rabbit (IgG) | 1:400 | 10mM Trisodium citrate (pH 6.0), 12 minutes microwave, 12 minutes cooling. | Abcam | Ab183322 (RRID:AB_2909463) |
| Tumour necrosis factor alpha (**TNF-α**) | Pro-inflammatory cytokine; increased expression by activated microglia, monocytes and some reactive astrocytes | Mouse (IgG1) | 1:500 | | Abcam | Ab1793 (RRID:AB_302615) |
| Translocator protein (**TSPO**) | Mitochondrial translocator protein; increased expression in inflammatory microglia and some reactive astrocytes | Rabbit (IgG) | 1:250 | | Abcam | Ab109497 (RRID:AB_10862345) |
| Ionised calcium-binding adaptor molecule 1 (**Iba-1**) | Microglia and monocytes | Rabbit (IgG) | 1:14,000 | | Proteintech | 10904-1-AP (RRID:AB_2224377) |
| C reactive protein (**CRP**) | Acute phase protein | Rabbit (IgG) | 1:800 | | Abcam | Ab32412 (RRID:AB_726934) |
| Osteopontin (**OP**) | Acute phase protein; secreted glycophosphoprotein | Rabbit (IgG) | 1:10,000 | 10mM Tris 1mM EDTA (pH 9.0), 40 minutes pressure cooker. | Proteintech | 22952-1-1AP (RRID:AB_2783651) |
| Alpha 1-antichymotrypsin; AACT (**Serpin A3**) | Acute phase glycoprotein | Mouse (IgG2b) | 1:25,000 | | Proteintech | 66078-1-Ig (RRID:AB_11182502) |
| Myeloperoxidase (**MPO**) | Abundant protein stored within granules of neutrophils | Rabbit (IgG) | 1:8000 | | Proteintech | 22225-1-AP (RRID:AB_2879037) |

Note: Anti-HLA-DR, anti-CRP and anti-osteopontin antibodies were sensitive to long formalin fixation durations, thus tissues which were fixed in formalin for more than 1 year were not included in immunohistochemistry experiments using these antibodies.

**Supplementary Table 6** Primary and secondary antibodies for multiplex immunofluorescence microglial oxidative phosphorylation assay

| Primary antibody | | | | | Secondary antibody | | | |
|---|---|---|---|---|---|---|---|---|
| **Target** | **Host (isotype)** | **Dilution** | **Antibody supplier** | **Catalogue number (RRID)** | **Alexa Fluor conjugate** | **Dilution** | **Antibody supplier** | **Catalogue number (RRID)** |
| Ionised calcium-binding adaptor molecule 1 (**Iba-1, microglial marker**) | Rabbit (IgG) | 1:1500 | Proteintech | 10904-1-AP (RRID:AB_2224377) | Alexa Fluor 405, anti-rabbit IgG | 1:100 | ThermoFisher Scientific | A31556 (RRID:AB_221605) |
| Cytochrome *c* oxidase subunit I (**COXI, Complex IV subunit**) | Mouse (IgG2a) | 1:200 | Abcam | Ab14705 (RRID:AB_2084810) | Alexa Fluor 488, anti-mouse IgG2a | 1:100 | ThermoFisher Scientific | A21131 (RRID:AB_2535771) |
| Voltage-dependent anion channel 1 (**VDAC1 / Porin, mitochondrial mass marker**) | Mouse (IgG2b) | 1:200 | Abcam | Ab14734 (RRID:AB_443084) | Alexa Fluor 546, anti-mouse IgG2b | 1:100 | ThermoFisher Scientific | A21143 (RRID:AB_2535779) |
| NADH:ubiquinone oxidoreductase subunit B8 (**NDUFB8, Complex I subunit**) | Mouse (IgG1) | 1:100 | Abcam | Ab110242 (RRID:AB_10859122) | XX Goat Biotin, anti-mouse IgG1 | 1:200 | ThermoFisher Scientific | A10519 (RRID:AB_1500809) |
| | | | | | Alexa Fluor 647, anti-streptavidin conjugate | 1:100 | ThermoFisher Scientific | S32357 |

Note: NDUFB8 primary antibody signal was amplified using an anti-mouse IgG1 biotinylated antibody and Alexa Fluor 647 anti-streptavidin antibody. 1mM EDTA (pH 8.0) was used for antigen retrieval. Long-fixed tissues were not included for immunofluorescence experiments.

**Supplementary Table 7: Top 20 significantly decreased proteins in the occipital cortex of patients with POLG-related epilepsy**. Rank of proteins (1 – 20) is determined by the lowest adjusted *P* value (Q value). The log 2 fold change (log2FC) for each protein in patients versus control occipital cortex tissues is also presented.

| Rank | Gene | Brief protein description | Occipital log2FC | Occipital Q value |
|---|---|---|---|---|
| **1** | LRPPRC | Leucine Rich Pentatricopeptide Repeat Containing protein; transcriptional regulator of nuclear and mitochondrial genes | -1.02 | 1.71E-13 |
| **2** | NDUFB3 | Mitochondrial complex I accessory subunit | -1.12 | 4.66E-13 |
| **3** | NDUFS5 | Mitochondrial complex I accessory subunit | -1.37 | 1.51E-12 |
| **4** | NDUFS1 | Mitochondrial complex I core subunit | -1.09 | 2.21E-12 |
| **5** | MT-ND4 | mtDNA-encoded complex I core subunit | -1.74 | 5.66E-12 |
| **6** | NDUFS6 | Mitochondrial complex I accessory subunit | -1.42 | 8.86E-12 |
| **7** | NDUFA10 | Mitochondrial complex I accessory subunit | -1.11 | 1.70E-11 |
| **8** | NDUFS2 | Mitochondrial complex I core subunit | -1.01 | 4.43E-11 |
| **9** | NDUFA9 | Mitochondrial complex I accessory subunit | -1.14 | 2.45E-10 |
| **10** | NDUFB9 | Mitochondrial complex I accessory subunit | -1.13 | 3.41E-10 |
| **11** | NDUFV1 | Mitochondrial complex I core subunit | -1.00 | 3.69E-10 |
| **12** | NDUFB7 | Mitochondrial complex I accessory subunit | -1.08 | 3.96E-10 |
| **13** | MRPL50 | Mitochondrial ribosomal protein L50 | -1.44 | 5.35E-10 |
| **14** | NDUFS3 | Mitochondrial complex I core subunit | -1.04 | 9.30E-10 |
| **15** | HCN1 | Hyperpolarization activated cyclic nucleotide gated potassium channel 1 | -1.37 | 1.12E-09 |
| **16** | GPR158 | G protein-coupled receptor 158; glycine metabotropic receptor | -1.13 | 2.29E-09 |
| **17** | SEC61A2 | SEC61 translocon subunit alpha 2; mediates the transport of polypeptides across the endoplasmic reticulum | -2.44 | 2.29E-09 |
| **18** | MT-ND5 | mtDNA-encoded complex I core subunit | -1.17 | 3.53E-09 |
| **19** | NDUFS8 | Mitochondrial complex I core subunit | -0.98 | 3.94E-09 |
| **20** | SMAP2 | Small ArfGAP2; stromal membrane-associated GTPase-activating protein 2 | -0.71 | 5.19E-09 |

Note: Complex I = NADH:Ubiquinone Oxidoreductase.

**Supplementary Table 8: Differential expression of the mitochondrial proteome**. 540 of 1136 MitoCarta 3.0 proteins were included for analysis in the proteomics study. Of the 540 analysed MitoCarta proteins, the number of differentially expressed MitoCarta proteins are presented for each comparison.

| **Protein list** | **No. of proteins** | **No. of MitoCarta proteins** | **No. of DEP MitoCarta proteins FC ± >1.5** |
|---|---|---|---|
| ***Total analysed proteins*** | ***N = 3790*** | ***N = 540*** | ***N/A*** |
| ANOVA *DEP* | N = 2196 | N = 321 / 540 | N/A |
| BA17 POLG vs BA17 Control *DEP* | N = 1225 | N = 227 / 540 | **N = 130 / 540** (18 up, 112 down) |
| BA9 POLG vs BA9 Control *DEP* | N = 363 | N = 74 / 540 | **N = 44 / 540** (6 up, 38 down) |
| BA17 Control vs BA9 Control *DEP* | N = 843 | N = 147 / 540 (134 up, 13 down) | **N = 35 / 450** (31 up, 4 down) |

***DEP*** Differentially expressed proteins.

*DEP* in the control occipital cortex versus control frontal cortex include n=53 Complex I, n=2 Complex II, n=3 Complex III, n=5 Complex IV, n=5 Complex V, n=2 electron carriers (OXPHOS n=70).

#### Supplementary Table 9: Significantly dysregulated pathways in POLG-related epilepsy patient tissues

| | Reactome pathway | Number of DEP | Adjusted *P* value | Cluster |
|---|---|---|---|---|
| **Decreased pathways in POLG patient occipital cortex** | | | | |
| 1 | Complex I Biogenesis | 36 | 1.63E-20 | Mitochondrial OXPHOS |
| 2 | Respiratory Electron Transport | 50 | 4.56E-15 | Mitochondrial OXPHOS |
| 3 | Mitochondrial Translation Initiation | 18 | 8.87E-11 | Mitochondrial translation |
| 4 | Mitochondrial Translation | 19 | 7.98E-10 | Mitochondrial translation |
| 5 | Mitochondrial Translation Elongation | 18 | 1.02E-09 | Mitochondrial translation |
| 6 | Mitochondrial Translation Termination | 17 | 1.27E-09 | Mitochondrial translation |
| 7 | Aerobic Respiration and Respiratory Electron Transport | 53 | 9.85E-08 | Mitochondrial OXPHOS |
| 8 | Neuronal System | 58 | 4.69E-07 | Neuronal proteins |
| 9 | Neurexins and Neuroligins | 19 | 5.35E-05 | Synaptic structure |
| 10 | Transmission Across Chemical Synapses | 42 | 2.57E-04 | Synaptic transmission |
| 11 | Protein-protein Interactions at Synapses | 22 | 2.57E-04 | Synaptic transmission |
| 12 | Unblocking of NMDA Receptors, Glutamate Binding and Activation | 11 | 6.98E-04 | Synaptic transmission |
| 13 | Neurotransmitter Receptors and Postsynaptic Signal Transmission | 30 | 7.59E-03 | Synaptic transmission |
| 14 | Transcriptional Regulation by MECP2 | 11 | 1.58E-02 | Gene expression |
| 15 | Long-term Potentiation | 9 | 1.67E-02 | Synaptic transmission |
| 16 | Negative Regulation of NMDA Receptor-Mediated Neuronal Transmission | 9 | 2.61E-02 | Synaptic transmission |
| 17 | Complex IV Assembly | 10 | 2.79E-02 | Mitochondrial OXPHOS |
| 18 | Synaptic Adhesion-Like Molecules | 8 | 3.07E-02 | Synaptic structure |
| 19 | Presynaptic Depolarization and Calcium Channel Opening | 6 | 4.20E-02 | Synaptic transmission |
| 20 | Activation of NMDA Receptors and Postsynaptic Events | 19 | 4.46E-02 | Synaptic transmission |
| **Decreased pathways in POLG patient frontal cortex** | | | | |
| 1 | Complex I Biogenesis | 18 | 2.44E-16 | Mitochondrial OXPHOS |
| 2 | Respiratory Electron Transport | 19 | 9.84E-11 | Mitochondrial OXPHOS |
| 3 | Mitochondrial Translation Initiation | 9 | 2.30E-08 | Mitochondrial translation |
| 4 | Mitochondrial Translation Termination | 9 | 2.30E-08 | Mitochondrial translation |
| 5 | Aerobic Respiration and Respiratory Electron Transport | 20 | 3.01E-08 | Mitochondrial OXPHOS |
| 6 | Mitochondrial Translation Elongation | 9 | 4.10E-08 | Mitochondrial translation |
| 7 | Mitochondrial Translation | 9 | 8.45E-08 | Mitochondrial translation |
| 8 | Translation | 12 | 2.72E-02 | Translation |
| 9 | Mitochondrial RNA Degradation | 3 | 2.72E-02 | Mitochondrial translation |
| 10 | Enhanced Binding of GP1BA Variant to VWF Multimer Collagen | 2 | 3.42E-02 | Haemostasis |
| 11 | Defective Binding of VWF Variant to GPIb IX V | 2 | 3.42E-02 | Haemostasis |
| 12 | Defects of Platelet Adhesion to Exposed Collagen | 2 | 3.42E-02 | Haemostasis |
| 13 | Cholesterol Biosynthesis | 3 | 3.57E-02 | Cholesterol metabolism |
| 14 | Activation of Gene Expression by SREBF (SREBP) | 3 | 4.76E-02 | Cholesterol metabolism |
| 15 | Anchoring Fibril Formation | 2 | 4.76E-02 | Extracellular matrix |
| 16 | Crosslinking of Collagen Fibrils | 2 | 4.76E-02 | Extracellular matrix |
| 17 | Diseases of Hemostasis | 2 | 4.76E-02 | Haemostasis |
| **Increased pathways in POLG patient occipital cortex** | | | | |
| 1 | Neutrophil Degranulation | 44 | 1.04E-12 | Innate immunity and inflammation |
| 2 | Innate Immune System | 56 | 4.01E-11 | Innate immunity and inflammation |
| 3 | Immune System | 68 | 4.22E-08 | Innate immunity and inflammation |
| 4 | Platelet Degranulation | 14 | 2.54E-04 | Platelets and haemostasis |
| 5 | Complement Cascade | 8 | 2.54E-04 | Complement system activation |
| 6 | Response to Elevated Platelet Cytosolic Ca2+ | 14 | 3.91E-04 | Platelets and haemostasis |
| 7 | Synthesis of Prostaglandins (PG) and Thromboxanes (TX) | 5 | 5.86E-04 | Eicosanoid metabolism and signalling |
| 8 | Initial Triggering of Complement | 6 | 9.61E-04 | Complement system activation |
| 9 | Regulation of Complement Cascade | 7 | 1.19E-03 | Complement system activation |
| 10 | Arachidonate Metabolism | 6 | 5.58E-03 | Eicosanoid metabolism and signalling |
| 11 | Platelet Activation, Signalling and Aggregation | 17 | 6.05E-03 | Platelets and haemostasis |
| 12 | Chaperone Mediated Autophagy | 5 | 9.17E-03 | Autophagy and degradation |
| 13 | Caspase-mediated Cleavage of Cytoskeletal Proteins | 4 | 9.17E-03 | Autophagy and degradation |
| 14 | Erythrocytes Take up Oxygen and Release Carbon Dioxide | 4 | 9.17E-03 | Gas transport in erythrocytes |
| 15 | Glycogen Breakdown (Glycogenolysis) | 4 | 1.64E-02 | Energy metabolism |
| 16 | Hemostasis | 25 | 1.73E-02 | Platelets and haemostasis |
| 17 | Glutathione Conjugation | 6 | 1.73E-02 | Detoxification |
| 18 | Dissolution of Fibrin Clot | 3 | 1.73E-02 | Platelets and haemostasis |
| 19 | Biological Oxidations | 10 | 1.85E-02 | Detoxification |
| 20 | O2 CO2 Exchange in Erythrocytes | 4 | 1.85E-02 | Gas transport in erythrocytes |
| 21 | Classical Antibody-Mediated Complement Activation | 4 | 1.85E-02 | Complement system activation |
| 22 | Creation of C4 and C2 Activators | 4 | 1.85E-02 | Complement system activation |
| 23 | Erythrocytes Take up Carbon Dioxide and Release Oxygen | 4 | 1.85E-02 | Gas transport in erythrocytes |
| 24 | Peptide Ligand-Binding Receptors | 4 | 2.83E-02 | Innate immunity and inflammation |
| 25 | Signaling by High-Kinase Activity BRAF Mutants | 6 | 4.06E-02 | MAPK and growth factor signalling |
| 26 | Interleukin-12 Family Signaling | 7 | 4.06E-02 | Innate immunity and inflammation |
| 27 | Apoptotic Cleavage of Cellular Proteins | 5 | 4.59E-02 | Autophagy and degradation |
| 28 | Antimicrobial Peptides | 3 | 4.59E-02 | Innate immunity and inflammation |
| 29 | RAF-independent MAPK1 3 Activation | 3 | 4.59E-02 | MAPK and growth factor signalling |
| 30 | Growth Hormone Receptor Signaling | 3 | 4.59E-02 | MAPK and growth factor signalling |
| 31 | Glycogen Metabolism | 4 | 4.59E-02 | Energy metabolism |

| | | | | |
|---|---|---|---|---|
| 32 | MAP2K and MAPK Activation | 6 | 4.59E-02 | MAPK and growth factor signalling |
| 33 | Passive Transport by Aquaporins | 2 | 4.59E-02 | Membrane transport |
| 34 | IFNG Signalling Activates MAPKs | 2 | 4.59E-02 | Innate immunity and inflammation |
| **Increased pathways in POLG patient frontal cortex** | | | | |
| 1 | Innate Immune System | 23 | 5.01E-04 | Innate immunity and inflammation |
| 2 | Immune System | 29 | 1.17E-03 | Innate immunity and inflammation |
| 3 | Complement Cascade | 5 | 4.46E-03 | Complement system activation |
| 4 | Regulation of Complement Cascade | 4 | 2.96E-02 | Complement system activation |
| 5 | Cellular Response to Heat Stress | 6 | 2.96E-02 | Heat shock response |
| 6 | Neutrophil Degranulation | 13 | 2.96E-02 | Innate immunity and inflammation |
| 7 | Regulation of HSF1-mediated Heat Shock Response | 5 | 2.96E-02 | Heat shock response |
| 8 | Classical Antibody-Mediated Complement Activation | 3 | 2.96E-02 | Complement system activation |
| 9 | Creation of C4 and C2 Activators | 3 | 2.96E-02 | Complement system activation |
| 10 | Gastrin-CREB Signalling Pathway via PKC and MAPK | 3 | 2.96E-02 | MAPK and growth factor signalling |

Pathway enrichment analysis was performed using ENRICHR and the Reactome database (accessed 06/11/2025). Analysis was conducted using significantly increased and decreased proteins (adjusted *P* value < 0.05 / 5.00E-02, and log2 fold change ± 0.58) against a custom background gene list composed of all analysed proteins (n = 3790). The number of differentially expressed proteins (**DEP**) used for pathway analyses included: n=201 increased and n=535 decreased proteins in POLG patient occipital cortex versus control occipital cortex, and n=77 increased and n=92 decreased proteins in POLG patient frontal cortex versus control frontal cortex. All significantly differentially expressed pathways are presented, based on an adjusted *P* value < 0.05. Altered Reactome pathways were grouped into functional clusters to identify shared functional changes.

**Supplementary Table 10: Top 20 decreased proteins in POLG-related epilepsy.** List of proteins with the greatest decreased log 2 fold change (log2FC) in occipital and frontal cortical tissues from patients with POLG-related epilepsy compared to respective control groups. Rank of proteins (1 – 20) is determined by the highest decreased log2FC in patient occipital cortex tissues. All proteins were significantly differentially expressed in both brain regions ($Q < 0.05$).

| | Gene | Brief protein description | Function overview | Occipital log2FC | Frontal log2FC | Occipital Q value | Frontal Q value |
|---|---|---|---|---|---|---|---|
| **1** | HMGCS1 | 3-Hydroxy-3-Methylglutaryl-CoA synthase 1 | Metabolism | -2.46 | -2.25 | 0.000016 | 0.000029 |
| **2** | SEC61A2 | Translocon subunit alpha 2 | Transporter protein | -2.44 | -0.88 | 0.000000 | 0.014091 |
| **3** | ACTL6B | Actin-like protein 6B | Actin-related protein | -2.37 | -2.01 | 0.002096 | 0.006684 |
| **4** | MRPL44 | Mitochondrial ribosomal protein L44 | Mitochondrial translation | -2.15 | -2.23 | 0.000955 | 0.000313 |
| **5** | RASGRF1 | Ras guanine nucleotide releasing factor | Guanine nucleotide exchange factor | -2.14 | -0.93 | 0.000001 | 0.030657 |
| **6** | BTBD17 | BTB domain containing 17 | Negative regulation of viral genome replication | -1.92 | -1.96 | 0.018044 | 0.010068 |
| **7** | MRPS27 | Mitochondrial ribosomal protein S27 | Mitochondrial translation | -1.91 | -1.04 | 0.000015 | 0.014852 |
| **8** | MRPL4 | Mitochondrial ribosomal protein L4 | Mitochondrial translation | -1.81 | -1.31 | 0.000000 | 0.000027 |
| **9** | MT-ND4 | Mitochondrially-encoded complex I subunit | Mitochondrial OXPHOS | -1.74 | -0.94 | 0.000000 | 0.000003 |
| **10** | MRPL38 | Mitochondrial ribosomal protein L38 | Mitochondrial translation | -1.67 | -1.41 | 0.000385 | 0.001727 |
| **11** | MYO1B | Myosin IB | Actin-related protein | -1.67 | -1.27 | 0.000620 | 0.006931 |
| **12** | ANKRD34A | Ankyrin repeat domain-containing protein | Mediates protein-protein interactions | -1.63 | -1.28 | 0.000167 | 0.001949 |
| **13** | HM13 | Signal peptide peptidase | Endoplasmic reticulum intramembrane proteolysis | -1.56 | -1.37 | 0.020531 | 0.036347 |
| **14** | NDUFA7 | Mitochondrial complex I accessory subunit | Mitochondrial OXPHOS | -1.53 | -0.72 | 0.000000 | 0.003556 |
| **15** | MRPL50 | Large mitochondrial ribosomal subunit | Mitochondrial translation | -1.44 | -0.73 | 0.000000 | 0.000160 |
| **16** | NDUFS6 | Mitochondrial complex I accessory subunit | Mitochondrial OXPHOS | -1.42 | -0.58 | 0.000000 | 0.000340 |
| **17** | DGKZ | Diacylglycerol kinase zeta | | -1.41 | -0.73 | 0.000001 | 0.005037 |
| **18** | MRPS34 | Small mitochondrial ribosomal subunit | Mitochondrial translation | -1.40 | -1.03 | 0.001253 | 0.015612 |
| **19** | UBLCP1 | Ubiquitin-like domain containing CTD phosphatase 1 | Proteasome regulation | -1.38 | -1.05 | 0.000772 | 0.008650 |
| **20** | HCN1 | Hyperpolarisation activated cyclic nucleotide gated potassium channel | Potassium channel | -1.37 | -0.46 | 0.000000 | 0.018457 |

**Supplementary Table 11: Top 20 increased proteins in POLG-related epilepsy.** List of proteins with the greatest increased log 2 fold change (log2FC) in occipital and frontal cortical tissues from patients with POLG-related epilepsy compared to respective control groups. Rank of proteins (1 – 20) is determined by the highest increased log2FC in patient occipital cortex tissues. All proteins were significantly differentially expressed in both brain regions ($Q < 0.05$).

| | Gene | Brief protein description | Function overview | Immune involvement | Occipital log2FC | Frontal log2FC | Occipital Q value | Frontal Q value |
|---|---|---|---|---|---|---|---|---|
| **1** | CRP | C-reactive protein | Acute phase response protein | Yes | 4.04 | 3.37 | 0.000063 | 0.000402 |
| **2** | SPP1 | Osteopontin, secreted phosphoprotein 1 | Cytokine | Yes | 3.91 | 2.34 | 0.000002 | 0.002330 |
| **3** | SCIN | Scinderin | Actin-binding protein | | 3.66 | 2.00 | 0.000034 | 0.021483 |
| **4** | HSPB8 | Heat shock protein B8, chaperone of BAG3 | Autophagy | | 3.33 | 2.26 | 0.000169 | 0.008439 |
| **5** | BAG3 | BAG family chaperone regulator 3, autophagy protein | Autophagy | | 3.17 | 1.74 | 0.000211 | 0.047020 |
| **6** | C1QB | Complement C1q subcomponent B | Complement initiation | Yes | 3.11 | 1.96 | 0.000001 | 0.000810 |
| **7** | CD14 | Monocyte differentiation factor | Toll-like receptor 4 co-receptor | Yes | 3.08 | 2.07 | 0.000000 | 0.000033 |
| **8** | HSPB6 | Heat shock protein beta-6 | Misfolded protein response | | 2.86 | 2.12 | 0.000002 | 0.000113 |
| **9** | TYMP | Thymidine phosphorylase | Nucleotide metabolism | | 2.82 | 1.58 | 0.000003 | 0.004997 |
| **10** | PIR | Pirin | Transcriptional coregulator of NFKβ | Yes | 2.82 | 1.55 | 0.000047 | 0.024282 |
| **11** | SERPINA3 | Alpha-1-anti-chymotrypsin, serine protease inhibitor | Acute phase response protein; inhibits cathepsin G | Yes | 2.80 | 2.47 | 0.000000 | 0.000000 |
| **12** | C1QC | Complement C1q subcomponent C | Complement initiation | Yes | 2.52 | 1.64 | 0.000000 | 0.000042 |
| **13** | RNASET2 | Ribonuclease T2 | Degrades RNA from microbial pathogens | Yes | 2.39 | 1.27 | 0.000117 | 0.044733 |
| **14** | HSPB1 | Heat shock protein B1, molecular chaperone | Misfolded protein response | | 2.19 | 1.76 | 0.000001 | 0.000013 |
| **15** | MGST1 | Microsomal glutathione S-transferase 1 | Anti-oxidant | Yes | 2.10 | 1.55 | 0.002874 | 0.026319 |
| **16** | CLU | Clusterin | Complement cytolysis inhibitor | Yes | 1.89 | 0.95 | 0.000000 | 0.000494 |
| **17** | CAPG | Capping actin protein | Actin regulatory protein | | 1.88 | 0.85 | 0.000000 | 0.014630 |
| **18** | CSF1R | Colony stimulating factor 1 receptor | Cytokine receptor | Yes | 1.81 | 1.00 | 0.000010 | 0.010528 |
| **19** | ORM1 | Orosomucoid 1 | Acute phase response protein | Yes | 1.75 | 1.99 | 0.003138 | 0.000379 |
| **20** | SERPINA1 | Alpha-1-anti-chymotrypsin, serine protease inhibitor | Acute phase response protein; Inhibits neutrophil elastase | Yes | 1.75 | 1.33 | 0.000000 | 0 |

### Supplementary Table 12: Enriched pathways in the control occipital cortex group.

| | Reactome pathway | Number of DEP | Adjusted P value | Cluster |
|---|---|---|---|---|
| **Increased pathways in Control occipital cortex** | | | | |
| 1 | Processing of Capped Intron-Containing Pre-mRNA | 60 | 1.36E-39 | RNA processing, splicing & maturation |
| 2 | mRNA Splicing | 54 | 9.35E-37 | RNA processing, splicing & maturation |
| 3 | mRNA Splicing - Major Pathway | 54 | 9.35E-37 | RNA processing, splicing & maturation |
| 4 | Metabolism of RNA | 75 | 1.65E-23 | RNA processing, splicing & maturation |
| 5 | mRNA Splicing - Minor Pathway | 17 | 1.83E-11 | RNA processing, splicing & maturation |
| 6 | SUMOylation | 20 | 1.83E-11 | SUMOylation & post-translational regulation |
| 7 | SUMO E3 Ligases SUMOylate Target Proteins | 18 | 8.56E-10 | SUMOylation & post-translational regulation |
| 8 | Transport of Mature mRNA Derived From an Intron-Containing Transcript | 13 | 5.91E-07 | mRNA transport & nuclear-cytoplasmic trafficking |
| 9 | Gene Expression (Transcription) | 47 | 7.96E-07 | Transcription & transcriptional elongation |
| 10 | Transport of Mature Transcript to Cytoplasm | 13 | 8.81E-07 | mRNA transport & nuclear-cytoplasmic trafficking |
| 11 | RNA Polymerase II Transcription Termination | 12 | 3.03E-06 | Transcription & transcriptional elongation |
| 12 | snRNP Assembly | 10 | 1.98E-05 | RNA processing, splicing & maturation |
| 13 | Metabolism of Non-Coding RNA | 10 | 1.98E-05 | RNA processing, splicing & maturation |
| 14 | SUMOylation of RNA Binding Proteins | 8 | 6.11E-05 | SUMOylation & post-translational regulation |
| 15 | FGFR2 Alternative Splicing | 10 | 8.16E-05 | Growth factor signalling & cellular differentiation |
| 16 | mRNA 3'-End Processing | 9 | 2.03E-04 | RNA processing, splicing & maturation |
| 17 | RNA Polymerase II Transcription | 37 | 3.39E-04 | Transcription & transcriptional elongation |
| 18 | Interactions of Vpr With Host Cellular Proteins | 8 | 3.71E-04 | Viral transcription, replication & host interaction |
| 19 | Viral Messenger RNA Synthesis | 12 | 3.71E-04 | Viral transcription, replication & host interaction |
| 20 | SUMOylation of DNA Replication Proteins | 7 | 7.14E-04 | SUMOylation & post-translational regulation |
| 21 | HIV Life Cycle | 13 | 1.29E-03 | Viral transcription, replication & host interaction |
| 22 | Transcriptional Regulation of Brown and Beige Adipocyte Differentiation | 5 | 1.76E-03 | Growth factor signalling & cellular differentiation |
| 23 | Transcriptional Regulation of Brown and Beige Adipocyte Differentiation by EBF2 | 5 | 1.76E-03 | Growth factor signalling & cellular differentiation |
| 24 | Positive Epigenetic Regulation of rRNA Expression | 7 | 3.24E-03 | Epigenetic & RNA-mediated regulation |
| 25 | tRNA Processing | 9 | 3.32E-03 | RNA processing, splicing & maturation |
| 26 | Epigenetic Regulation of Gene Expression | 10 | 3.32E-03 | Epigenetic & RNA-mediated regulation |
| 27 | SUMOylation of Chromatin Organization Proteins | 6 | 3.32E-03 | SUMOylation & post-translational regulation |
| 28 | Transport of Mature mRNA Derived From an Intronless Transcript | 6 | 3.32E-03 | mRNA transport & nuclear-cytoplasmic trafficking |
| 29 | Transport of Mature mRNAs Derived From Intronless Transcripts | 6 | 3.32E-03 | mRNA transport & nuclear-cytoplasmic trafficking |
| 30 | Transport of the SLBP Dependant Mature mRNA | 6 | 3.32E-03 | mRNA transport & nuclear-cytoplasmic trafficking |
| 31 | Transport of the SLBP Independent Mature mRNA | 6 | 3.32E-03 | mRNA transport & nuclear-cytoplasmic trafficking |
| 32 | SUMOylation of DNA Damage Response and Repair Proteins | 7 | 3.86E-03 | SUMOylation & post-translational regulation |
| 33 | RNA Polymerase II Transcribes snRNA Genes | 4 | 4.13E-03 | RNA processing, splicing & maturation |
| 34 | Nuclear Import of Rev Protein | 6 | 5.11E-03 | Viral transcription, replication & host interaction |
| 35 | Vpr-mediated Nuclear Import of PICs | 6 | 5.11E-03 | Viral transcription, replication & host interaction |
| 36 | Transcriptional Regulation by Small RNAs | 7 | 5.25E-03 | Epigenetic & RNA-mediated regulation |
| 37 | Defective TPR May Confer Susceptibility Towards Thyroid Papillary Carcinoma (TPC) | 5 | 5.48E-03 | Cancer-associated processes |
| 38 | Processing of Capped Intronless Pre-mRNA | 5 | 5.48E-03 | RNA processing, splicing & maturation |
| 39 | Regulation of Glucokinase by Glucokinase Regulatory Protein | 5 | 5.48E-03 | Growth factor signalling & cellular differentiation |
| 40 | HCMV Early Events | 10 | 6.97E-03 | Viral transcription, replication & host interaction |
| 41 | Adipogenesis | 6 | 6.98E-03 | Growth factor signalling & cellular differentiation |
| 42 | Rev-mediated Nuclear Export of HIV RNA | 6 | 6.98E-03 | Viral transcription, replication & host interaction |
| 43 | SLBP Dependent Processing of Replication-Dependent Histone Pre-mRNAs | 4 | 8.71E-03 | RNA processing, splicing & maturation |
| 44 | SLBP Independent Processing of Histone Pre-mRNAs | 4 | 8.71E-03 | RNA processing, splicing & maturation |
| 45 | Nuclear Pore Complex (NPC) Disassembly | 5 | 8.71E-03 | mRNA transport & nuclear-cytoplasmic trafficking |
| 46 | SUMOylation of SUMOylation Proteins | 5 | 8.71E-03 | SUMOylation & post-translational regulation |
| 47 | Signaling by FGFR2 | 8 | 9.11E-03 | Growth factor signalling & cellular differentiation |
| 48 | Late Phase of HIV Life Cycle | 10 | 1.30E-02 | Viral transcription, replication & host interaction |
| 49 | tRNA Processing in the Nucleus | 6 | 1.30E-02 | mRNA transport & nuclear-cytoplasmic trafficking |
| 50 | Interactions of Rev With Host Cellular Proteins | 6 | 1.30E-02 | Viral transcription, replication & host interaction |
| 51 | NEP NS2 Interacts With the Cellular Export Machinery | 5 | 1.30E-02 | Viral transcription, replication & host interaction |
| 52 | SUMOylation of Ubiquitinylation Proteins | 5 | 1.30E-02 | SUMOylation & post-translational regulation |
| 53 | Transport of Ribonucleoproteins into the Host Nucleus | 5 | 1.30E-02 | RNA processing, splicing & maturation |
| 54 | HCMV Infection | 11 | 1.74E-02 | Viral transcription, replication & host interaction |
| 55 | Signaling by FGFR | 8 | 1.74E-02 | Growth factor signalling & cellular differentiation |
| 56 | Gene Silencing by RNA | 7 | 1.86E-02 | Epigenetic & RNA-mediated regulation |
| 57 | Export of Viral Ribonucleoproteins From Nucleus | 5 | 1.94E-02 | Viral transcription, replication & host interaction |
| 58 | Abortive Elongation of HIV-1 Transcript in the Absence of Tat | 3 | 2.32E-02 | Viral transcription, replication & host interaction |
| 59 | Formation of the Early Elongation Complex | 3 | 2.32E-02 | Transcription & transcriptional elongation |
| 60 | Formation of the HIV-1 Early Elongation Complex | 3 | 2.32E-02 | Viral transcription, replication & host interaction |
| 61 | IRF3-mediated Induction of Type I IFN | 3 | 2.32E-02 | Innate immune & anti-viral signalling |
| 62 | STING Mediated Induction of Host Immune Responses | 3 | 2.32E-02 | Innate immune & anti-viral signalling |
| 63 | Formation of HIV Elongation Complex in the Absence of HIV Tat | 4 | 2.36E-02 | Viral transcription, replication & host interaction |
| 64 | Formation of HIV-1 Elongation Complex Containing HIV-1 Tat | 4 | 2.36E-02 | Viral transcription, replication & host interaction |
| 65 | HIV Transcription Elongation | 4 | 2.36E-02 | Viral transcription, replication & host interaction |
| 66 | SUMOylation of Transcription Cofactors | 4 | 2.36E-02 | SUMOylation & post-translational regulation |
| 67 | Tat-mediated Elongation of the HIV-1 Transcript | 4 | 2.36E-02 | Viral transcription, replication & host interaction |

| | | | | |
|---|---|---|---|---|
| 68 | NS1 Mediated Effects on Host Pathways | 5 | 3.62E-02 | Viral transcription, replication & host interaction |
| 69 | Formation of RNA Pol II Elongation Complex | 4 | 3.83E-02 | Transcription & transcriptional elongation |
| 70 | RNA Polymerase II Transcription Elongation | 4 | 3.83E-02 | Transcription & transcriptional elongation |
| 71 | HCMV Late Events | 7 | 4.07E-02 | Viral transcription, replication & host interaction |
| 72 | 2-LTR Circle Formation | 3 | 4.47E-02 | Viral transcription, replication & host interaction |
| 73 | mRNA Capping | 3 | 4.47E-02 | RNA processing, splicing & maturation |
| 74 | FGFR2 Mutant Receptor Activation | 3 | 4.47E-02 | Growth factor signalling & cellular differentiation |
| 75 | RNA Polymerase I Transcription Initiation | 3 | 4.47E-02 | Transcription & transcriptional elongation |
| 76 | Signalling by FGFR2 IIIa TM | 3 | 4.47E-02 | Growth factor signalling & cellular differentiation |
| 77 | Regulation of Endogenous Retroelements | 5 | 4.50E-02 | Epigenetic & RNA-mediated regulation |

Pathway enrichment analysis was performed using ENRICHR and the Reactome database (accessed 06/11/2025). Analysis was conducted using significantly increased and decreased proteins (adjusted $P$ value < 0.05 / 5.00E-02, and log2 fold change ± 0.58) in control occipital cortex versus control frontal cortex, against a custom background gene list composed of all analysed proteins (n = 3790). The number of differentially expressed proteins (**DEP**) used for pathway analyses included: n=293 significantly increased proteins in control occipital cortex and n=105 significantly decreased proteins in control occipital cortex. Significantly increased pathways in control occipital cortex are presented, based on an adjusted $P$ value < 0.05. No significantly decreased pathways in control occipital cortex were identified (adjusted $P > 0.05$). Altered Reactome pathways were grouped into functional clusters to identify shared functional changes.

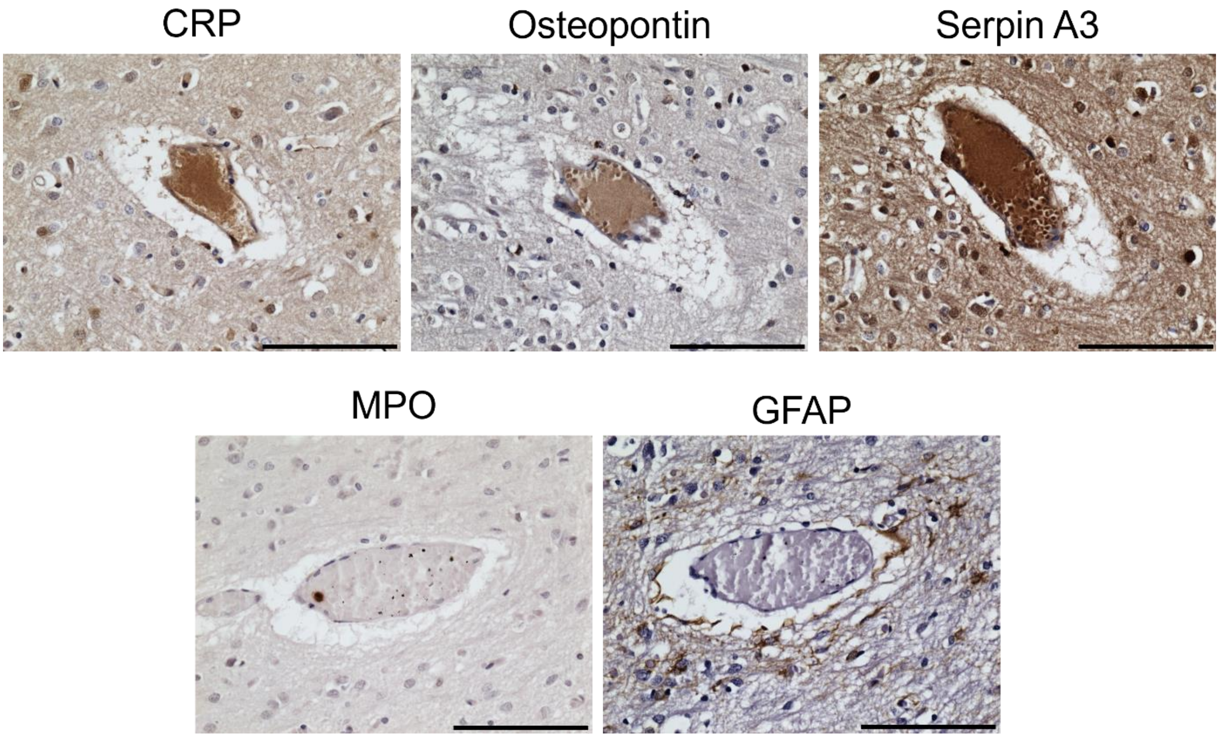


**Supplementary Fig. 1 Increased immunoreactivity of inflammatory proteins in vessels in POLG-related epilepsy**. *CRP* C-reactive protein; *MPO* Myeloperoxidase; *GFAP* Glial fibrillary acidic protein. Patient 11 Occipital cortex. Scale bar = 100 µm

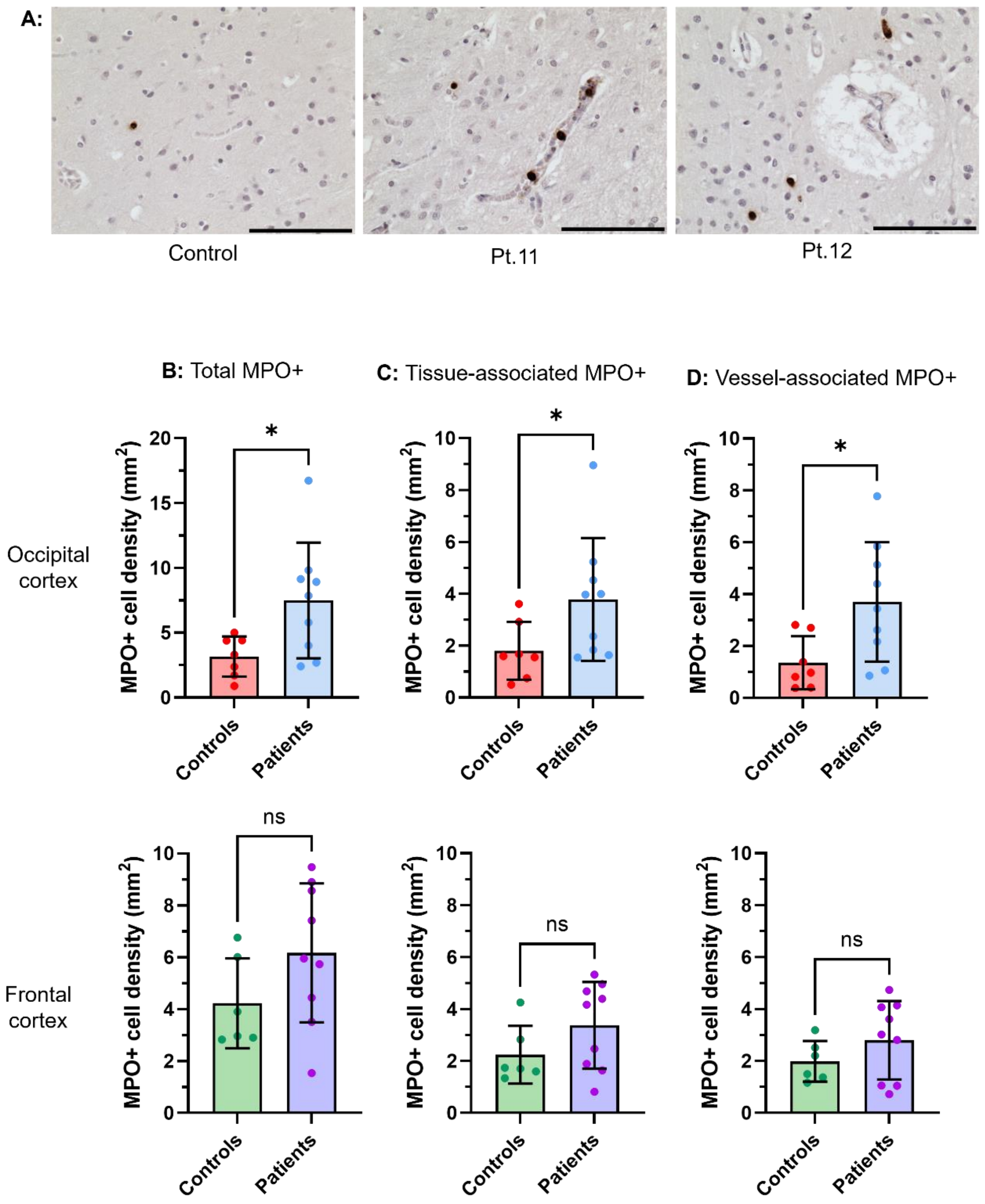


**Supplementary Fig. 2 Increased myeloperoxidase protein expression in the occipital cortex of patients with POLG-related epilepsy.**

(**A**) Representative images demonstrate myeloperoxidase (MPO)-immunoreactive (+) cells in control and patient occipital cortex tissues. In the control tissue, the example MPO+ cell appears to be localised within the brain parenchyma. In the patient tissues, MPO+ cells appear to be localised both within the brain parenchyma and within blood vessels. Tissue is counterstained with Mayer's Haematoxylin. Scale bars = 100 µm.

The total density of MPO+ cells (**B**), in addition to the density of MPO+ cells that appear to be localised within the brain parenchyma (**C**: tissue-associated MPO+) and vasculature (**D**: vessel-associated MPO+) are presented for control and patient tissues in the occipital and frontal cortex. Individual circles indicate individual cases. Error bars = standard deviation. Data analysed using Mann-Whitney u test. * = $P < 0.05$; ns = non-significant.
